# *NF2* lacking exon 11 induced by antisense gene therapy is able to partially recover merlin deficiency on *NF2*-SWN iPSC-derived spheroid model

**DOI:** 10.1101/2025.11.14.688164

**Authors:** Gemma Casals-Sendra, Ignasi Jarne-Sanz, Núria Catasús, Irene Boluda-Luis, Ariadna Quer, Emilio Amilibia, Conxi Lázaro, Eduard Serra, Ignacio Blanco, Elisabeth Castellanos, NF-SWN Spanish Reference Center (CSUR on Phakomatosis)

## Abstract

*NF2*-related Schwannomatosis (*NF2*-SWN) is an inherited autosomal dominant disorder resulting from loss-of-function mutations in the *NF2* gene, for which no effective treatment is currently available. Furthermore, truncating variants in *NF2* are associated with the severest phenotype compared to in-frame or missense variants.

Previously, a shorter *NF2* isoform with exon 11 skipped (merlin_e-11), induced through antisense morpholino oligomers (PMOs), was able to partially rescue the deleterious effect of nonsense variants located at that exon in patients’ primary fibroblast. To test the potential of this approach into Schwann cells, those responsible of *NF2*-SWN tumours, we developed an iPSC-based model carrying heterozygous and homozygous truncating variants on exon 11 of the *NF2* gene and differentiated them to Schwann cells forming spheroids. After 3 days of treatment, merlin_e-11 expression was induced in NF2-deficient cell lines. Furthermore, key pathways associated with NF2-deficiency in schwannomas, such as PI3K/Akt/mTORC and YAP levels, were recovered without signs of toxicity. These results confirm that the PMO treatment induces effective skipping of exon 11 in Schwann cell spheroids, generating an hypomorphic merlin_e-11 that has the capacity to partially rescue merlin-deficiency in *NF2*-SWN spheroid cell model and it being a potential treatment of patients harbouring truncating variants located in exon 11.

## INTRODUCTION

*NF2*-related Schwannomatosis (*NF2*-SWN) previously called Neurofibromatosis type 2, is an autosomal dominant (AD) disease caused by loss-of-function (LOF) variants in the *NF2* gene (22q12.2), which encodes for merlin protein (moesin-ezrin-radixin like) (1). *NF2*-SWN patients present a predisposition to the development of multiple benign tumours (schwannomas, meningiomas and ependymomas) affecting both the central and peripheral nervous system, in addition to cutaneous and ocular manifestations (2,3) among others. The hallmark of the disease is the development of bilateral vestibular schwannomas (VS) (4).

Despite being benign, the development of these tumours is associated with a significant increase in morbidity, primarily due to their multiplicity and anatomically sensitive locations. This phenomenon is often accompanied by symptoms such as sensorineural hearing loss in 90% of patients, multifocal weakness, pain and immobility (5,6). Without treatment, the potential for tumours to be life-threatening is contingent upon their location. Consequently, *NF2*-SWN reduces both patient quality of life and life expectancy (4,7). Conventionally, *NF2*-SWN patients have been offered tumour management, either through surgery or stereotactic radiosurgery (8). However, the multifocal presentation of *NF2*-SWN tumours represents significant therapeutic challenges (9). Accordingly, different therapeutic approaches that target merlin-related pathways are being tested.

Merlin contributes to cellular cytoskeletal organization (10); activates the Hippo pathway, which regulates YAP/TAZ targets (11); and inhibits mTORC1 signalling (12). Finally, it has been demonstrated that merlin also participates in the regulation of TP53, VEGF and epithelial-to-mesenchymal transition signalling, all of which are known to be deregulated in schwannomas (13–19).

Currently, Bevacizumab (20), an anti-angiogenic agent, Everolimus (21), an mTOR inhibitor, or Brigatinib, an inhibitor of ALK and multiple other tyrosine kinases (22), have shown radiographic responses in multiple tumour types and improvement in hearing function. However, due to their limited efficacy for some *NF2*-SWN related tumours (23) and long-term treatment significant adverse effects (9,24) there remains a great need for the development of patient-centred therapies that go beyond tumour-directed strategies.

In this context, the strong genotype-phenotype correlation that enables prediction of *NF2*-SWN evolution constitutes an opportunity to modulate *NF2*-SWN disease by directly modifying merlin function. *NF2* truncating variants are associated with a more severe phenotype, whereas missense mutations and small in-frame deletions tend to correspond to milder phenotypes with a lower tumour burden (25,26). In accordance with this, a strategy based on Antisense Oligomers (ASOs) (27), specifically Phosphorodiamidate Morpholino Oligomers (PMOs), was tested in primary fibroblasts derived from *NF2*-SWN patients to induce the skipping of exon 11 of the *NF2* gene. This approach was able to induce an hypomorphic merlin_e-11 protein while exhibiting partial recovery of functionality and a rescue of the cytoskeleton phenotype (28).

Among the various non-viral therapeutic strategies that have been employed, antisense gene therapy using ASOs has demonstrated notable success in modulating gene expression and treating various hereditary diseases. ASOs are characterized as single-stranded and have been observed to act at the RNA level. Specifically, they have been shown to correct or modify the expression of the target protein, leading to the production of a less deleterious form (29,30). As an example, Eteplirsen, a FDA approved ASO therapy has been demonstrated to induce milder forms of DMD in Duchenne patients by the induction of skipping of DMD gene exon 51 (31,32). Other examples, transmitted in an autosomal recessive pattern include spinal muscular atrophy (SMA) (33), homozygous familial hypercholesterolemia (HoFH) (34). Moreover, there is evidence of autosomal dominant diseases, including retinitis pigmentosa (RP) (35), amyotrophic lateral sclerosis (36) or Huntington’s disease (37).

Since *NF2*-SWN schwannomas arise due to a secondary *NF2* mutation in Schwann cells (SCs), which exhibit somatic full merlin deficiency (38,39), it will be desirable to assess the effectiveness of the ASO-based strategy in rescuing merlin-deficient Schwann cells, by using PMOs, the same strategy has been applied for DMD patients. However, preclinical *in vitro* models to study *NF2*-SWN and to develop new therapies show relevant limitations: primary SCs cultures derived from VS are perishable, resulting in a limited lifespan (40,41), while immortalized VS-derived or meningioma-derived cell lines (42–44) do not fully replicate the genetic and pathophysiological features of benign *NF2*-related tumours (41).

To circumvent the limitation, we generated a novel pair of induced pluripotent stem cell (iPSC) lines harbouring a truncating variant at exon 11 in one or both alleles of the *NF2* gene. iPSCs retain the genomic background from the cell of origin and can be differentiated towards the neural crest and Schwann cell axis (45) *in vitro*. This results in cells that exhibit an expression profile closely resembling that of human merlin-deficient SCs. Consequently, they are considered a valuable *in vitro* model system for studying the pathogenesis of *NF2*-SWN Schwann cells and for evaluating new therapeutic approaches (45,46). In this study, a model for exon 11 was stablished: *NF2*(+/-) and *NF2*(-/-) SC were cultured under 3D conditions (SC-like spheroids) and treated with PMOs, the ASOs utilized in primary fibroblasts (28), resulting in the successful induction of the merlin_e-11 hypomorphic protein. The therapeutic potential of this antisense gene therapy approach in rescuing *NF2*-deficient Schwann cells that harbour truncating variants in exon 11 was evaluated after a week of treatment.

## RESULTS

### Induced pluripotent stem cells harbouring single or bi-allelic LOF variants in the *NF2* gene at exon 11

With the aim of achieving iPSCs with truncating variants at exon 11 both in heterozygous and homozygous forms, we used the CRISPR/Cas9 technology to specifically edit exon 11 of *NF2* gene in a control *NF2*(+/+) iPSC line (FiPS Ctrl 1-SV4F-7). We obtained single-cell isogenic *NF2*(+/-) and *NF2*(-/-) iPSC clones (named 2H9 and 2F6, respectively). We fully characterized induced mutations through cloning the *NF2* coding region and Sanger sequencing to confirm compound heterozygous clones (Fig. S1A, Tab. S1).

Both *NF2*(+/-) and *NF2*(-/-) iPSC lines expressed cell surface proteins and transcription factors associated with pluripotency: Oct4, SSEA-3, SSEA-4, Nanog, Sox2, Tra-1-60 and Tra-1-81 (Fig. 1A). These were positive for alkaline phosphatase staining and showed karyotype stability after at least 20 passages (46, XY) (Fig. 1B and C). As previously described (45), *NF2*(-/-) iPSCs formed less compact colonies and underwent major spontaneous differentiation compared to control iPSCs, consistent with the known role of merlin in maintaining pluripotency (Fig. 1D) (45). Nevertheless, both *NF2*(+/-) and *NF2*(-/-) iPSCs showed the capacity to differentiate into the three primary germ layers *in vitro* through embryoid bodies (EBs) formation (Fig. 1E). In addition, all *NF2* genotypes were confirmed through a merlin western blot (Fig. 1F).

**Fig. 1.**
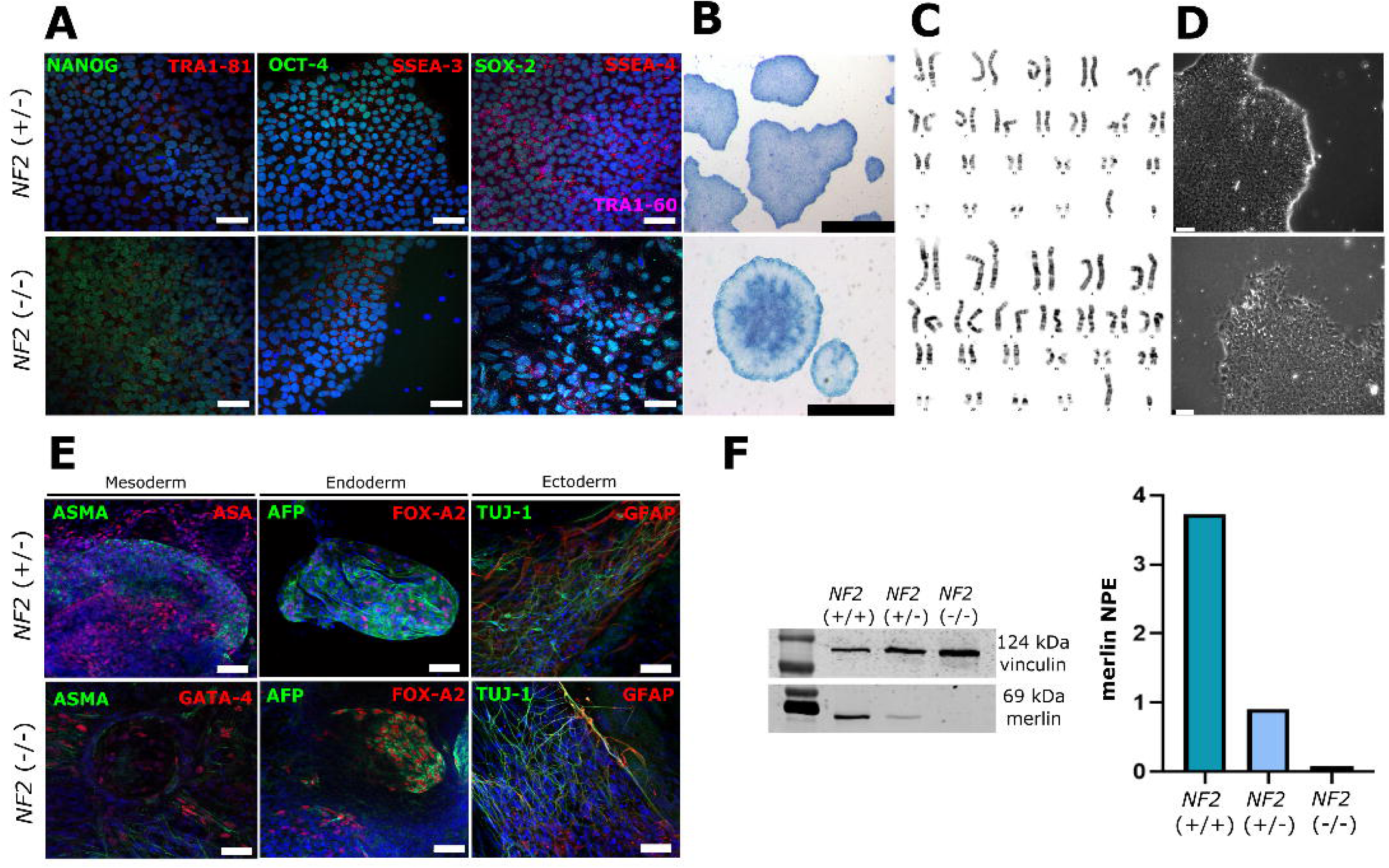
Characterization of iPSC Clones. **A:** Immunohistochemical analysis of pluripotency markers: NANOG, OCT4, and SOX2 (green); TRA-1-81, SSEA-3, and SSEA-4 (red); TRA-1-60 (pink). Cell nuclei were stained with DAPI (blue). Scale bar: 50 µm. **B:** Alkaline Phosphatase (ALP) staining of iPSC colonies. Scale bar: 1000 µm. **C:** Karyotype of iPSC lines at passage 20 (46, XY). **D:** Morphological assessment of iPSC colonies. Scale bar: 75 µm. **E:** Immunohistochemical analysis demonstrating the potential of differentiation into the three primary germ layers of the iPSC lines *in vitro*: mesoderm (ASMA, green; ASA, red for *NF2*(+/-); GATA-4, red for *NF2*(-/-)), endoderm (AFP, green; FOXA2, red), and ectoderm (TUJ1, green; GFAP, red). Scale bar: 50 µm. **F:** Merlin expression assessed by Western blot analysis. The FiPs line derived from fibroblasts *NF2*(+/+) was used as a control.

Genomic characterization of *NF2*(+/-) and *NF2*(-/-) iPSC lines by massive SNP-array genotyping and Whole Exome Sequencing (WES) showed no relevant differences with respect to the iPSC line of origin, except for the variants induced at *NF2* gene, confirming no significant off-target effect due to CRISPR-Cas9 editing (Fig. S1B, Tab. S2).

### *NF2* iPSC differentiation towards the NC-SC lineage

Given that the cells that initiate schwannoma formation in *NF2*-related SWN are *NF2*(-/-) cells derived from the SC lineage, we applied a differentiation protocol towards the NC-SC axis (Fig. 2A). After ten days of NC differentiation, cells achieved NC morphology (Fig. 2B). Cell identity was assessed by flow cytometry, immunocytochemistry and transcriptome analysis of NC markers (Fig. 2C, D and H). As expected from previous results (45), *NF2(+/-)* cells already showed high expression of p75 and Hnk1 after early passages of the NC differentiation process (92%), while *NF2*(-/-) cells showed lower expression of both markers at early passages (8.9%), but acquired higher percentages of p75 and Hnk1-positive cells at the mature NC stage (68,9%), compared with their control counterparts (Fig. 2D). Additionally, all lines expressed NC markers Sox10, L1CAM and p75 (NGFR) (Fig. 2H). However, some expression of S100B, a SC classical marker, was observed in *NF2(-/-)* NC immunocytochemical analysis, which was consistent with the heterogeneity detected by flow cytometry and as previously described (Fig. 2C) (45). Neither *NF2*(+/-) or *NF2*(-/-) NC cells showed altered capacity of migration, but proliferation rates were lower when compared to control NC cells (Fig. 2E and F). As a last step, NC cells were cultured in SC differentiation medium (SCDM) for 14 days in 3D conditions, obtaining spheroids that stained positive for p75 and S100B (Fig. 2G).

**Fig 2.**
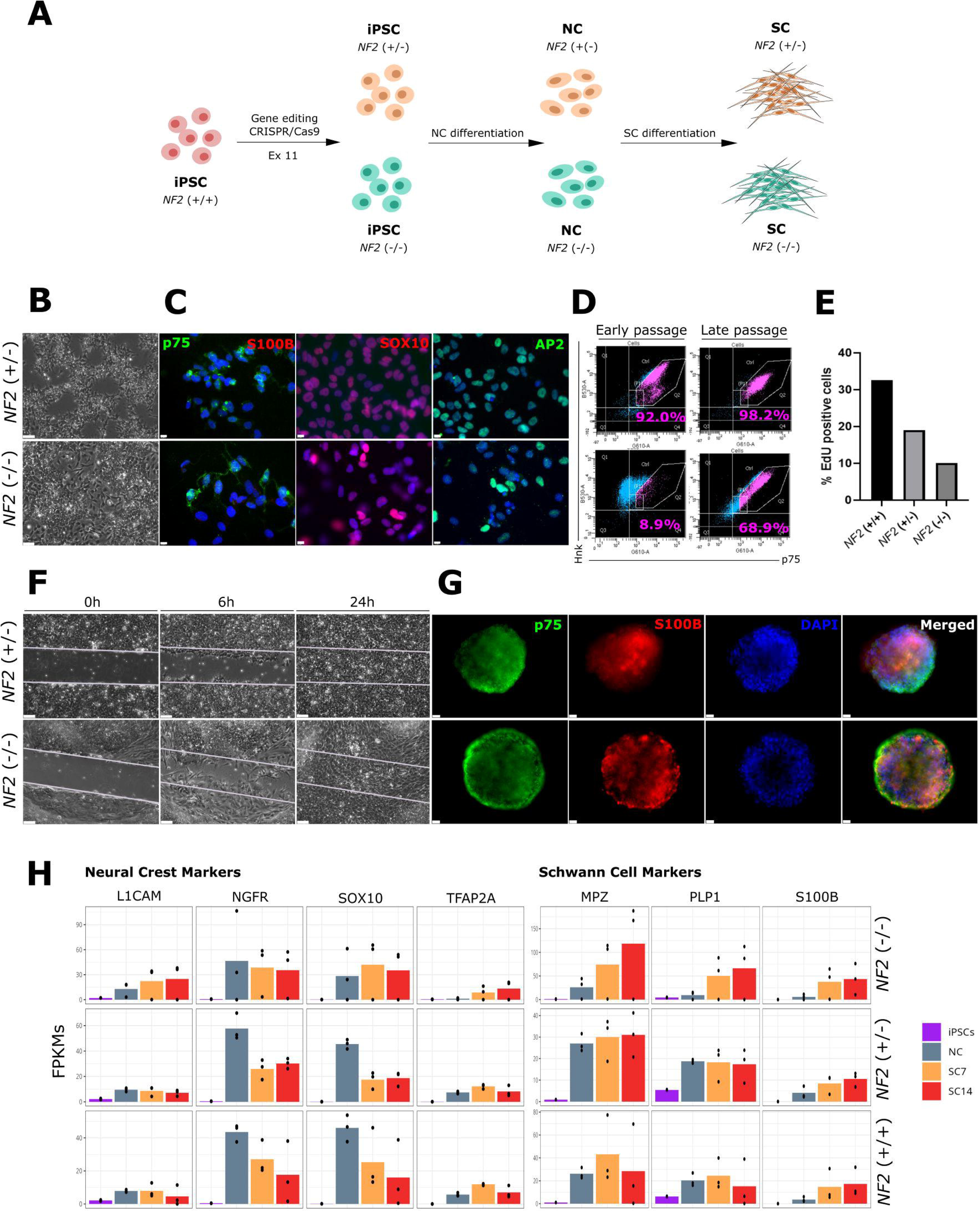
Differentiation of iPSC clones towards the Neural Crest – Schwann Cell lineage. **A:** Schematic representation of the experimental procedure applied for obtaining merlin-deficient Schwann Cell-like spheroids. **B:** Morphological assessment of Neural Crest (NC) cells in *NF2* (+/-) and *NF2* (-/-) cell lines. Scale bar: 75 µm. **C:** Immunocytochemical analysis of NC lineage markers: p75 (green), AP2 (green), S100B (red), and Sox10 (red). Cell nuclei were stained with DAPI (blue). Scale bar: 10 µm. **D:** Flow cytometry analysis of NC markers p75 and Hnk1. The percentage of p75 and Hnk1-positive cells is indicated in pink. Early and late passages correspond to passage 4 and passage 8, respectively. **E:** Proliferation analysis of the clones at the Neural Crest stage using Click-IT EdU assay. **F:** Scratch assay for the study of cell migration. Scale bar, 75µm. **G:** Immunohistochemical staining of p75 (green) and S100B (red) after 14 days of Schwann Cell differentiation. Cell nuclei were stained with DAPI (blue). Scale bar: 25 µm. **H:** Gene expression (FPKMs: Fragments Per Kilobase of transcript per Million mapped reads) of the classical Neural Crest (L1CAM, NGFR, SOX10 and TFAP2A) and Schwann Cell (MPZ, PLP1 S100B) markers of the *NF2*(-/-), *NF2*(+/-) and *NF2*(+/+) generated lines at the stage of iPSC, NC, SC at 7 days and 14 days of differentiation respectively.

To confirm these newly generated lines differentiate towards SC as has been previously described (45,47), we studied the transcriptome at iPSC, NC and SC at two time points of the differentiation process (7 and 14 days). We observed that iPSC, NC and SC triplicates clustered according to cell type through principal component analysis (PCA) but it is worth noting that *NF2*(-/-) NC show remarkable variability and tended to cluster with SCs, which is consistent with previous observations of a tendency to spontaneously differentiate, probably due to the lack of *NF2* (Fig. S2A). At the RNA level, both *NF2*(+/-) and *NF2*(-/-) cell lines expressed high levels of SC markers at 14 days of differentiation in a similar way to the control line, indicating that NC cells progress from a precursor state into a committed SC identity (Fig. 2H).

Furthermore, we confirmed that very few genes were differentially expressed (DEG) when comparing *NF2*(+/+) and *NF2*(+/-) expression profile (n=497) while *NF2*(-/-) SC-spheroids showed a higher number of DEGs (n=2017 and n=1358) when compared with *NF2*(+/-) and *NF2*(+/+) respectively. Gene Set Enrichment Analysis (GSEA) and Single Sample GSEA showed enrichment of pathways directly or indirectly regulated by merlin in Schwann cells and schwannomas such a PI3K/AKT/mTOR, Hippo NFKβ, p53, Wnt, and IL6-JAK-STAT3 (Fig. S2B and C). These results demonstrate that these new *NF2*(+/-) and *NF2*(-/-) SC spheroids harbouring truncating variants at exon 11 reproduce the phenotype and the expression profile of *NF2*(+/-) and *NF2*(-/-) SC spheroids already described (45).

We observed that other relevant merlin targets previously found to be altered in schwannomas were also upregulated; such as inflammatory response, epithelial-to-mesenchymal transition (EMT) or estrogenic response (Fig. S2B). These analyses indicated that *NF2*(-/-) SC-like spheroid could also recapitulate some profile signatures observed in *NF2*-related schwannomas. To investigate the potential of these *NF2*(-/-) SC spheroids to generate schwannomas, we performed an *in vivo* experiment and injected 2.25 million *NF2*(+/+), *NF2*(+/-) or *NF2*(-/-) SC forming spheroids at the 14 day differentiation time point into contact with the sciatic nerve of nude mice (Tab. S3, Fig. S3A). Four months after injection, none of the injected cells were able to generate a visible engrafted cell mass. Furthermore, after euthanasia of the mice, no schwannomas or other types of neural lesions were identified in the histological sections of *NF2*-deficient SC at 14 days of differentiation. The multiple neural bundles observed adjacent to skeletal muscle, varying in calibre and in some cases slightly branched, showed no remarkable morphological alterations. Likewise, we did not detect abnormalities in the other tissues examined, including skin, adipose tissue, cartilage, and lymph nodes (Fig. S3B). In spite of these results, we cannot discard the possibility of *NF2*(-/-) SC potential to generate schwannoma-like tumours. In this experiment, we were not able to generate tumours; however, further refinement of the technical conditions may be required to establish orthotopic models for benign tumours. Nevertheless, upon consideration of cellular phenotype and transcriptomic analyses, the developed spheroid model represents a promising platform for testing potential therapies for *NF2*-related schwannomatosis, such as the use of PMOs. This assertion is made in the light of the inherent limitations of *in vitro* culture, namely the absence of a tumour environment and the presence of cellular heterogeneity observed in schwannomas.

### Induction of skipping of merlin exon 11 through PMOs in iPSC and SC-spheroids

In previous studies, we have shown that a specific pair of PMOs complementary to both 5’- and 3’- intron-exon boundaries of exon 11 of the *NF2* gene (PMO_ES11) induce the in-frame deletion of that exon, producing a hypomorphic merlin, called merlin_e-11, in *NF2*-SWN primary fibroblasts (28). Here, we evaluated the therapeutic potential of this pair of PMOs (PMO_ES11) on iPSC and SC spheroids.

Firstly, we confirmed the efficacy of the PMO treatment at RNA level by time course studies on control *NF2*(+/+) iPSC, which showed skipping of full exon 11 at 40µM at 24h, 48h and 72h, achieving more than 50% of the exon-less form after 72h of treatment (Fig. 3A and B). As performed in primary fibroblast 2D cultures, we delivered PMOs through an endocytosis-mediated process with the use of the Endo-Porter molecule. Similar results were obtained when *NF2*(+/+) NC derived from control iPSC at 72h were treated (Fig. 3B).

**Fig 3:**
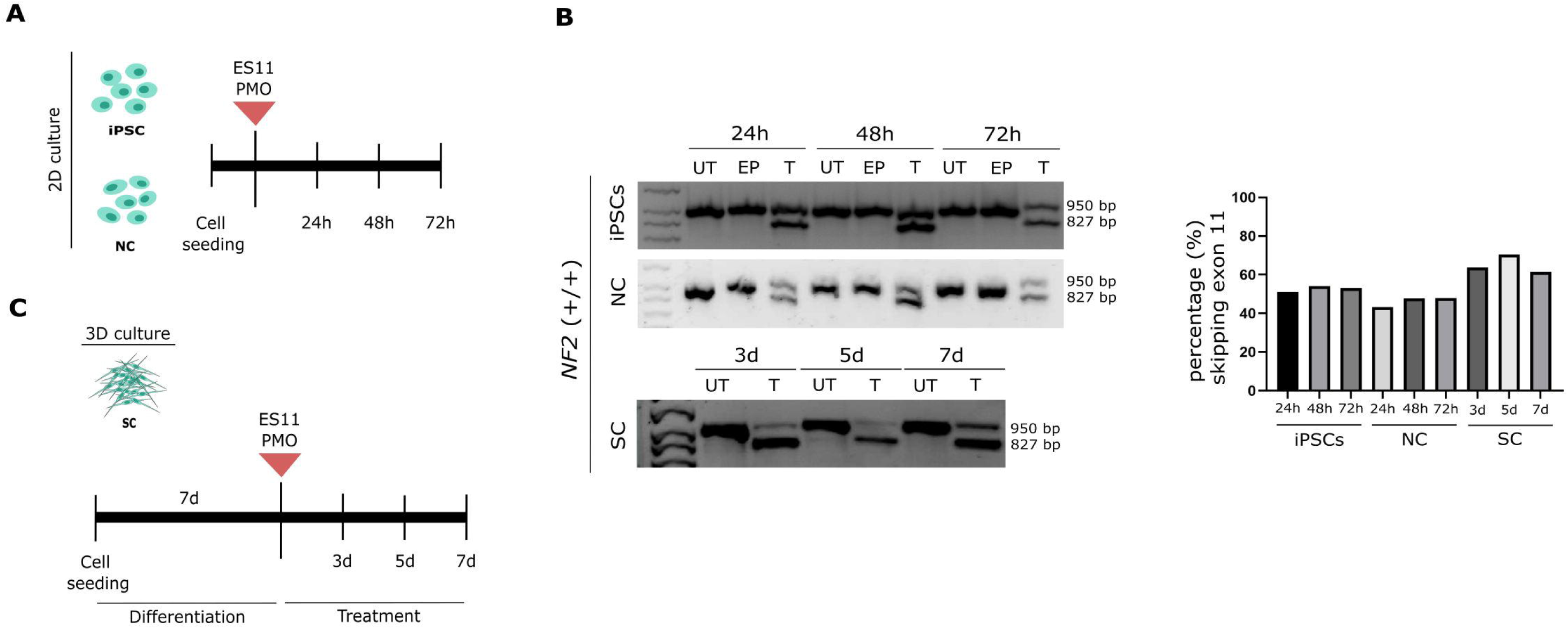
Induction of skipping of merlin exon 11 through PMOs in iPSC and SC-spheroids. **A:** Schematic representation of the experimental planning applied for the treatment of 2D cultured cells (NC and iPSCs). **B:** Electrophoresis gel reveals the skipping of exon 11 of the treated transcripts in the *NF2*(+/+) treated cells. Bands are shown at 950bp and 827bp. UT stands for untreated cells, EP refers to the cells treated only with the vehicle, T stands for treated cells. **C:** Schematic representation of the experimental planning applied for the treatment of 3D cultured cells (SC-spheroids).

However, due to the half-life of these molecules and the fact that *NF2*(-/-) SC-like differentiated from iPSC only growth on 3D culture (SC-like spheroids), we set up the conditions to treat spheroids with a new pair of PMOs (PMO_ES11_vivo) which were targeting the same 5’- and 3’-intron-exon boundaries of exons 11. These PMO_ES11_vivo were chemically modified to be able to enter cells directly without the need of any delivery method (vivo-PMOs, GeneTools). In particular, the *in vivo* delivery moiety is an octoguanidine dendrimer which conjugates to the 3’ end of a Morpholino oligo to enhance cell penetration.

Control *NF2*(+/+) SC spheroids cultured for 14 days on SC-DIFF media, were treated at 2µM for 3, 5 and 7 days with PMO_ES11_vivo pair, obtaining approximately 70% of the exon-less form after 3 days of treatment at RNA level (Fig. 3B and C), achieving even better results than primary fibroblast and iPSC or NC cells without relevant signs of toxicity.

These results indicate for the first time that PMO therapeutic molecules can be effective both in 2D iPSC cultures and in 3D cultures derived from these cells, such as SC spheroids, resulting in exon 11 skipping; and that they have the potential to generate a hypomorphic merlin_e-11.

### Merlin_e-11 hypomorphic protein rescued the *NF2*-related alterations in *NF2*(-/-) SC spheroids

To investigate whether merlin_e-11 induction can restore the phenotype resulting from *NF2* deficiency in SC-like spheroids, we treated *NF2*(+/-) and *NF2*(-/-) SC spheroids as follows: NC were differentiated towards SC in SC-DIFF media, and premature SC were treated at day 7 with 2µM PMO_ES11_vivo pair for 3, 5 and 7 days, in a total of 14 days of culture under differentiation conditions (Fig. 3C). *NF2*(+/-) cells showed a similar proportion of exon 11 skipped as the control *NF2*(+/+) SC-spheroids (80’3%) (Fig. 4A and B). The merlin levels of this *NF2*(+/-) cells nearly reached the levels seen in control cells, probably due to the nonsense mediated decay (NMD) cellular mechanism degrading the *NF2* mRNAs harbouring the truncating variants in untreated cells (Fig. 4C). Similarly, *NF2*(-/-) expressed the shorter isoform in a predominant manner (89’6%) (Fig. 4A and B) which was corroborated at protein level (Fig. 4C) However, the amount of merlin does not seem to reach the levels of *NF2* (+/-) cells (Fig. 4C).

**Fig 4:**
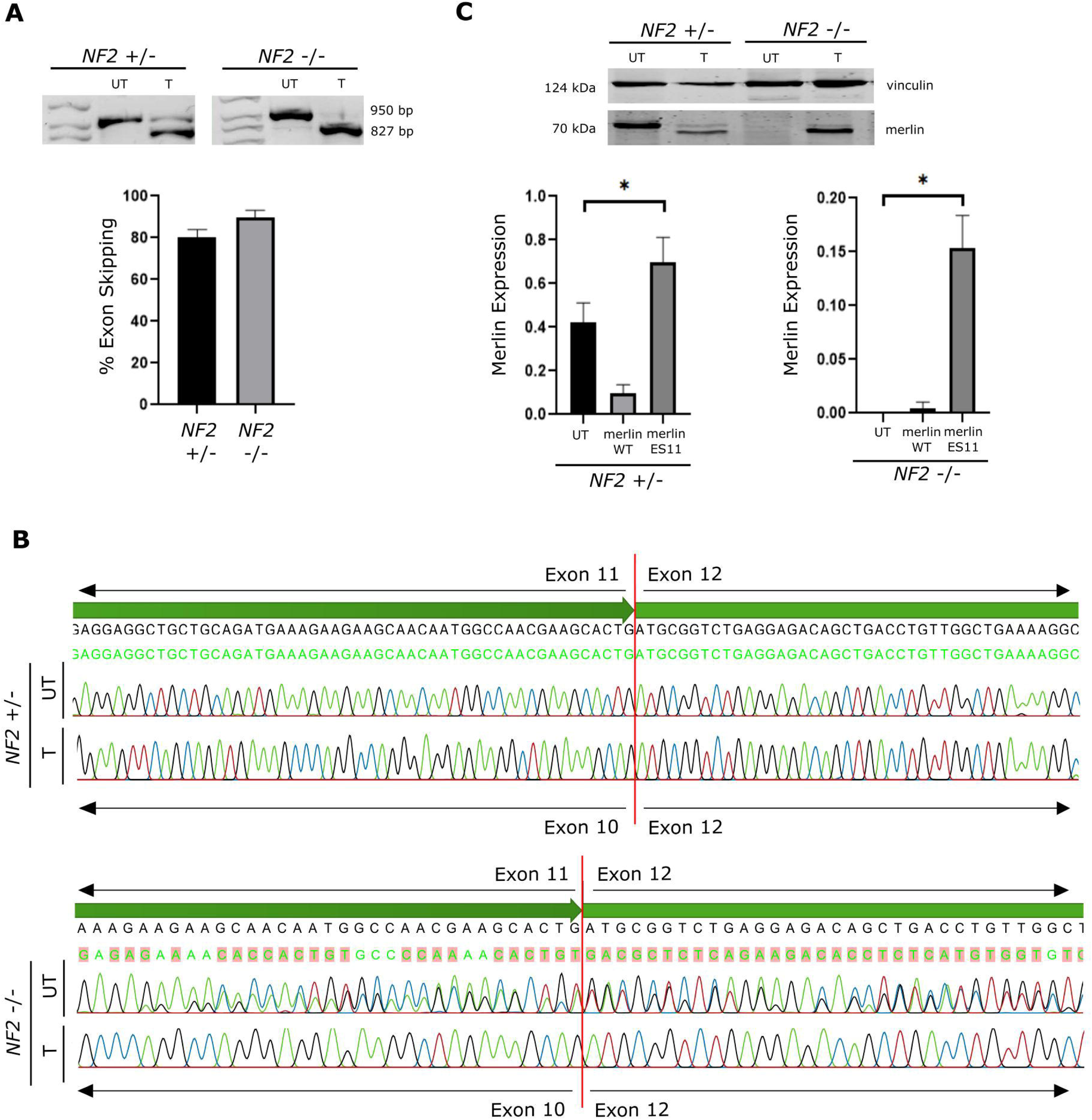
Generation of skipping of exon 11 and produce merlin protein. **A:** Electrophoresis gel reveals the absence of exon 11 in 80.3% and 89.6% of the treated transcripts in the *NF2*(+/-) and *NF2*(-/-) treated SC spheroids respectively. Bands in electrophoresis gel are shown at 950bp and 827bp. **B:** Sanger sequencing confirms the presence of the exon skipping were the presence of exon 12 is consecutive to exon 10 after the administration of the Vivo-Morpholino dose in both *NF2*(+/-) and *NF2*(-/-) SC spheroids. UT refers to the not treated cells. **C:** Merlin expression assessed by Western blot analysis. UT stands for untreated cells. Merlin WT refers to the merlin isoforms that do not present the exon skipping of exon 11. Merlin ES11 stands for the merlin that presents the exon skipping caused by the administration of the PMO treatment. Bars express mean ±SD levels of merlin expression from three independent experiments. Mann-Whitney U statistical test was performed among groups. Significant comparisons are shown as *<0.05; p=0.0213 [*NF2*(+/-)], p=0.0351 [*NF2*(-/-)].

Despite the low merlin levels induced on *NF2*(–/–), we decided to continue characterizing the effect of this merlin_e-11 on SC spheroids to determine whether these levels would be sufficient to rescue the merlin deficiency phenotype. We performed a differential expression analysis from bulk RNA analysis, comparing cells of the two disease-associated genotypes [*NF2* (+/-) and *NF2* (-/-)] differentiated for seven days and treated for seven additional days. Cells were treated with *NF2* exon 11-targeted PMO (PMO_ES11_vivo) or control PMO (scrambled). Firstly, to detect unspecific PMO responses, we identified the DEGs comparing untreated cells and scramble-treated cells. These were cross-referenced with the DEGs identified when comparing untreated cells with PMO_ES11_vivo-treated cells. We conducted an analysis for each tested genotype. This analysis resulted in the identification of 493 and 246 unspecific DEGs, respectively, for *NF2* (+/-) and *NF2* (-/-) genotypes (Fig. S4, Tab. S4). Then, we studied the interaction of these unspecific DEGs with *NF2* by STRING, and the predicted interaction between *SLC9A3R1* and *NF2* was the sole exception on *NF2*(+/-) cells, while none unspecific DEG identified in *NF2*(-/-) cells were found to interact with *NF2*, suggesting that the treatment of PMOs may induce a non-relevant off-target effect (Fig. S5 A).

Afterwards, we identified the number of specific DEGs on the induction of merlin_e-11 by PMOs (n =1,743 and n=2,738 in *NF2* (-/-) and *NF2* (+/-) cells, respectively) (Fig. S4A). Secondly, we conducted a functional enrichment analysis to explore trends induced by the treatment. As expected, very few pathways were significantly altered in *NF2*(+/-) and *NF2*(-/-) SC-like spheroids cells upon treatment. GO analysis highlights downregulation of genes related to cellular junction, cytoskeleton organization and neuron projection, while cytokine regulation and immune response were upregulated (Fig. S6). Furthermore, GSEA Hallmark signalling pathway analysis revealed that most *NF2*-related pathways that were significantly enriched (FDR > 0.05) in *NF2*(-/-) SC-like spheroids without treatment, changed trend after treatment. Notably, the mTORC1, G2M checkpoint, DNA repair, MYC targets, and E2F targets hallmarks, which were downregulated in *NF2*-deficient cells, revert their expression levels (ie. become upregulated) after seven days of PMO_ES11_vivo treatment compared to *NF2*(-/-) untreated (Fig. 5A1). Similarly, PMO treatment was able to reduce the aberrant upregulation of the EMT-related genes and protein secretion. Additionally, despite the few DEGs in *NF2*(+/-) SC-like spheroids, genes related to the G2M_checkpoint, mitotic spindle, E2F target and UV response hallmark pathways changed their expression profile after PMO treatment on *NF2*(+/-) SC-like spheroids (Fig. 5A2).

**Fig 5:**
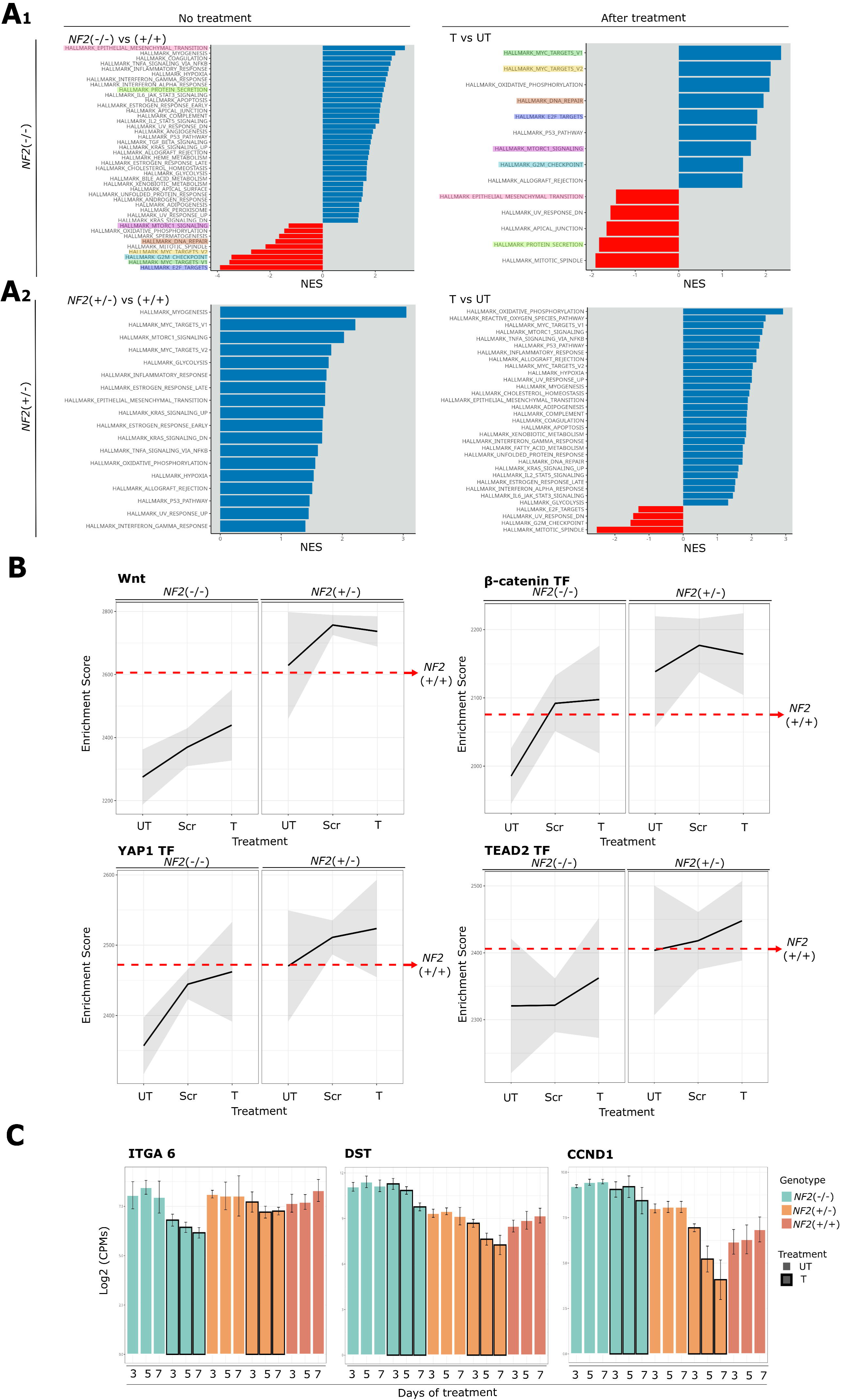
Expression profile of *NF2*-deficient lines after PMO treatment. **A:** GSEA of Schwann Cell spheroids after 7 days of PMO treatment. NES stands for Normalized Enrichment Score. Upregulated pathways represented in blue, downregulated pathways represented in red. Pathways were filtered by p value (< 0.05). Colour-marked hallmark pathways refer to those that recovered after treatment. **A1:** Comparison of *NF2*(-/-) vs. *NF2*(+/+) spheroids without treatment [left] and *NF2*(-/-) treated cells vs. *NF2*(-/-) untreated cells [right]. **A2:** Comparison of *NF2*(+/-) vs. *NF2*(+/+) spheroids without treatment [left] and *NF2*(+/-) treated cells vs. *NF2*(+/-) untreated cells [right]. **B:** Single Sample GSEA comparison. Treatment stands for untreated (UT), scrambled (Scr) and treated (T) Schwann Cell spheroids. Red arrow references to WT levels of expression. TF means Transcription Factor. **C:** Gene expression (log2(CPMs)) of NF2-related markers (ITGA6, DST and CCND1) of *NF2*(+/-) and *NF2*(-/-) after 3, 5 and 7 days of treatment. FiPs *NF2*(+/+) were used as a control. UT stands for untreated and T stands for treated cells.

To further investigate if these changes in the expression lead to a functional improvement, we studied the expression of genes that are described as being affected by the absence of merlin in primary or immortalised Schwann cells by single sample gene set enrichment analysis (ssGSEA) (Fig. 5B). We examined the levels of YAP and TEAD target genes, since their expression is negatively regulated by merlin. As expected from merlin deficiency, Hippo signalling was downregulated while TAZ target genes were upregulated in *NF2*(-/-) SC-like spheroids, whereas seven days after *NF2* exon 11-targeted PMO treatment and the induction of merlin_e-11, levels of YAP1-related TF were almost the same as *NF2*(+/+) levels, while TEAD2-related TF increased slightly but did not recover control levels. Similar results were observed for b-catenin/CTNNB1-related TF and Wnt target genes (Fig. 5B). Moreover, we investigated the normalised expression levels of selected genes described as being altered in schwannomas. Similarly to hallmark GSEA or ssGSEA shown results, we observed a clear trend toward wild type levels of cyclin D (*CCND1*) at day 3 of treatment, while the effect on DST was visible at day 7. For the laminin receptors α6 integrin (*Itga6*), a slight tendency of recovery could be observed at day 7, although it is not statistically significant due to the high interexperiment variability (Fig. 5C).

Finally, we were able to confirm that the restoration of the expression profile of most of *NF2*-related pathways had an impact at cellular level. Levels of cyclin D1 (*CCND1*) were also rescued in terms of the protein, which were accompanied by a reduction of pS6/s6 ratio and nuclear YAP, which suggests a reduction in cell cycle progression, and an inhibition of mTORC and Hippo pathway, all of them upregulated in *NF2*-deficient Schwann cells (Fig. 6A and B). A tendency of pAKT/AKT ratio recovery was also observed, even though it was not statistically significant after 7 days of treatment (Fig. 6A). We also observed a reduction of spheroid size after treatment, which is not associated with an increase of apoptosis, indicating that the induced merlin_e-11 could generate a reduction in cell size, as previously described (Fig. 6C and D)(48).

**Fig 6:**
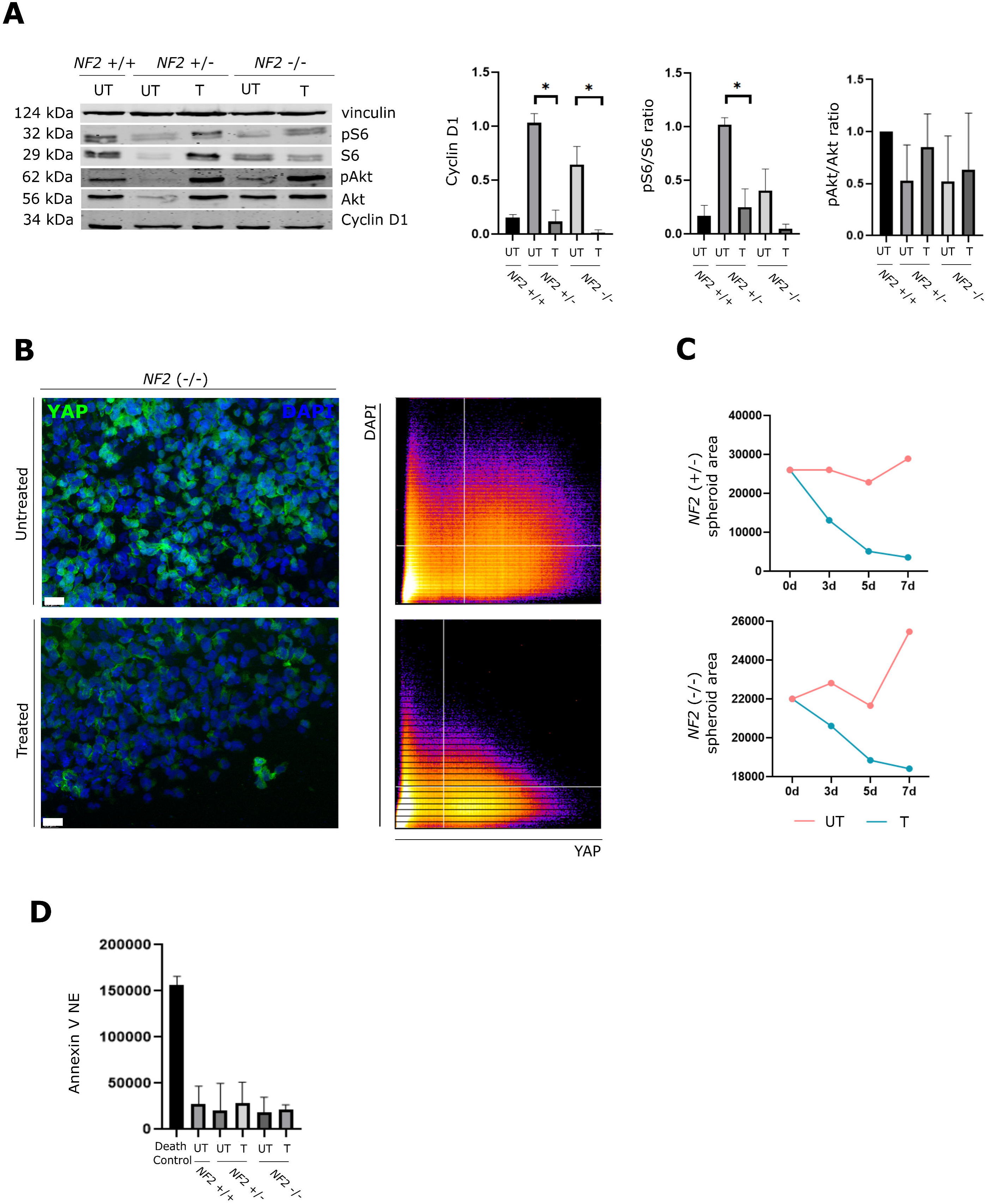
Functional assays confirm the partial rescue of *NF2*-SWN phenotype. **A:** Expression of *NF2*-related pathways in SC spheroids after treatment assessed by Western Blot analysis. The FiPs line derived from fibroblasts *NF2* (+/+) was used as a control. UT stands for untreated, T stands for treated. Bars express mean ±SD levels of merlin expression from three independent experiments. Levels of expression were normalized with the control, following by a min-max normalization. Mann-Whitney U statistical test was performed among groups. Significant comparisons are shown as *<0.05; p=0.0004 [CyclinD1 for *NF2*(+/-)], p=0.0208 [CyclinD1 for *NF2*(-/-)], p=0.0089 [pS6/S6 ratio for *NF2*(+/-)]. **B:** Immunohistochemical analysis demonstrating the levels of nuclear YAP in *NF2*(-/-) SC spheroids: YAP1 (green). Cell nuclei were stained with DAPI (blue). Scale bar: 25 µm. Cytofluorogram show correlation between green channel (YAP) and blue channel (DAPI). Area overlapped corresponds to 16% (Treated) and 6.32% (Untreated). **C:** Area quantification of spheroids untreated (red) and after 7 days of treatment (blue). **D:** Levels of apoptosis of SC spheroids after treatment were quantified by Annexin V assay. NE stands for Normalized Expression. UT stands for untreated cells, T stands for treated cells. A death control of cells induced apoptosis was used.

Altogether, these findings showed specific PMOs targeting exon 11 (PMO_ES11) can generate a shorter merlin isoform (merlin_e-11) in iPSC-derived SC spheroids, which is able to restore the expression profile and the function of the key *NF2*-related pathways altered in cells that originate schwannoma development, indicating that PMO_ES11_vivo induces a restoration of merlin function in *NF2*-deficient SC-like cells after 7 days of treatment. No apoptosis or other relevant cellular alterations were observed in *NF2*(+/-) SC due to the induction of two hypomorphic merlin instead of one wild-type and one defective merlin.

## DISCUSSION

*NF2*-related Schwannomatosis (*NF2*-SWN) is a multisystem genetic disorder for which the development of effective therapeutic options with no adverse consequences is needed. The clinical presentation of the disease is variable and correlated to the type of germline variant inherited in the *NF2* gene (25,26). In this context, our lab showed that the skipping of exon 11 of the *NF2* gene induced by the use of PMOs, produced an hypomorphic merlin_e-11 protein exhibiting partial recovery of merlin deficiency in primary fibroblasts derived from *NF2*-SWN patients (28). Despite these encouraging findings, preclinical *NF2*-SWN *in vitro* models, to test the potential of ASOs therapy available at that moment, primary SCs and immortalized tumor-derived cell lines, showed important limitations (40–44) and not fully replicate the genetic and pathophysiological features of benign *NF2*-SWN related tumors (41).

The aim of the present study was to determine if PMO-induced therapy can effectively induce the expression of merlin_e-11 and rescue the alterations due to merlin deficiency in a cellular model that mimics the different *NF2* genotypes present in *NF2*-SWN patients. Therefore, we tested the efficacy of ASO treatment in a new generated isogenic pair of iPSC that can be differentiated towards SCs (45), harbouring CRISPR/Cas9-induced truncating variants at *NF2* exon 11. These cells can resemble both *NF2*-SWN patient germline (*NF2* (+/-)) and schwannoma cell-of-origin genotype (*NF2* (-/-)). Our group has previously established an iPSC model with single or bi-allelic inactivation of *NF2*. This was achieved by combining the direct reprogramming of human primary vestibular schwannoma cells with the use of CRISPR/Cas9 *NF2* gene editing. This process resulted in the differentiation of the cells towards the Schwann cell lineage and the modeling of *NF2*-SWN *in vitro* (45). In the present study, to overcome the lack of iPSC model with truncating variants at exon 11, we generated a new *NF2*(+/-) and *NF2*(-/-) isogenic pair of iPSCs harbouring truncating variants on exon 11 by editing our control iPSC line. We also performed a functional comparison with the *NF2*(+/-) and *NF2*(-/-) iPSC lines already generated directly from schwannoma cells that definitely confirms that the described cellular alterations can be attributed to the absence of *NF2*.

Furthermore, our findings demonstrate that both iPSC and SC-like cells, when cultured in 3D conditions (spheroids), exhibit sensitivity to PMOs targeting exon 11 by effectively inducing the merlin_e-11 hypomorphic protein in both cell types. Furthermore, following only a week of treatment, it was possible to rescue the expression of most of the *NF2*-related pathways, such as mTORC, to reduce the levels of cyclin D1 and reduce the expression of nuclear YAP levels on *NF2*(-/-) SC-like spheroids. This finding indicates that, despite the limited duration of treatment, the induced PMO merlin_e-11 effectively restores the expression profile and the function of the key *NF2*-related pathways that are altered on schwannoma cell-of-origin. Notably, there is an absence of evidence of apoptosis or other significant cellular alterations.

Additionally, ASOs are known to broadly target cell types of brain *in vivo* mouse models (49,50) and iPSC-derived *in vitro* 2D cell cultures such as neurons and astrocytes (51,52). Our findings demonstrate the effectiveness of *in vivo* PMOs on 3D cell cultures, with no evidence of visible toxicity. This suggests that the efficacy of such therapies can be assessed in 3D *in vitro* models using the same molecules used in animal models, thereby accelerating their evaluation and facilitating their clinical application. This finding is significant for the future of ASO-based therapeutic development, particularly in the context of hereditary diseases. The potential establishment of models utilizing spheroids or organoids could be a valuable approach (53–55).

Despite the results of the *NF2*-SWN 3D SC model, there are some limitations that must be considered, including maintenance of SC commitment over long periods of time and the absence of cellular interaction and signaling provided by their physical interaction and proximity with other cell types. These cell types include neurons and other glial cells present in the nerves where schwannomas arise, as well as immune cell infiltration identified in schwannomas). This, in conjunction with the absence of trophic factors and a niche effect, could provide a rationale for the observed lack of schwannoma formation after engraftment on orthotopic model.

Furthermore, a pivotal issue concerning PMO merlin-e11 therapy for *NF2*-SWN patients pertains to the impact of transforming *NF2*(+/-) SC into SC expressing two hypomorphic merlin. Our findings on fibroblast cells suggest that the substance is well tolerated by patient cells (28). No significant alterations were observed in the bulk RNA-seq analysis or in the functional assays conducted on *NF2*(+/-) SC-like spheroids. However, it is imperative to note that additional *in vivo* analyses are necessary to definitively confirm the absence of long-term side effects.

Finally, the tested dose in this study is higher than FDA approved PMOs for other inherited diseases (56–58). Thus, redesigns and modifications of the tested PMO_ES11 antisense molecule should be performed to improve its efficacy. In addition, further pharmacodynamics and pharmacokinetics studies should be conducted to determine the appropriate range of doses and the associated toxicity and effectiveness. Additionally, the effectiveness of PMO_ES11 treatment should be evaluated through *in vivo* studies to ascertain whether the induction of hypomorphic merlin_e-11 would prevent tumour formation or reduce tumour burden when administered systemically or directly to tumours.

In summary, this study demonstrates that PMO-induced skipping of exon 11 could restore merlin function in an *NF2* iPSC model. It represents a critical next step in evaluating the potential medical approach for antisense therapy for *NF2*-SWN patients harboring germline truncating variants at exon 11 of the *NF2* gene. However, future work should focus on the effect of the PMO treatment systemically, and its therapeutic approach on tumour formation.

## MATERIAL AND METHODS

Detailed information will be found in Supplementary Materials and Methods.

### SAMPLES

All procedures performed were in accordance with the ethical standards of the IGTP Institutional Review Board (PI-24-044), which approved this study and with the 1964 Helsinki declaration and its later amendments.

### CELL CULTURE

#### iPSCs culture

iPSCs were cultured as described in (45). When necessary, differentiated cells were removed manually. In this study, the *NF2*(+/+) FiPS line (FiPS Ctrl 1-SV4F-7, (FiPS Ctrl 1-SV4F-7 registered in the Spanish National Stem Cell Bank/ ESiO44C in https://hpscreg.eu/)) was used in all experiments as a control.

#### Neural Crest differentiation

iPSC lines were differentiated to Neural Crest as previously described (59) Medium was changed every day, cells were maintained at 37°C and 5% CO2. Cell Culture conditions are described in Catasús, et al.2025 (45).

#### Schwann Cell differentiation in 3D

NC cells were detached with Accutase, 2.25·10^6^ cells/well were seeded onto AggreWell TM800 24-well plates (Stem Cell Technologies) in 2mL SCDM (described below) and cultured at 37°C and 5% CO2 as described in Catasús, et al.2025 (45). On days 7 and 14 spheroids were collected and processed for subsequent analysis. Cells were cultured for a maximum of 14 days.

### IPSC EDITING AND GENOMICS CHARACTERIZATION

#### CRISPR/Cas9 gene edition in iPSC lines

CRISPR/Cas9 editing was conducted using the ArciTect ribonucleoprotein (RNP) system (STEMCELL Technologies). The sgRNA targeted exon 11 of the *NF2* gene (TCGCTCGAGAGAAGCAGATG) and was designed using the Synthego CRISPR Design Tool (CRISPR Design Tool, 2024, v.1.3). Transfection was performed with the TransIT-X2® Dynamic Delivery System (Mirus) following the manufacturer’s instructions.

#### NF2 gene characterization after CRISPR/Cas gene editing

*NF2* exon 11 was screened by Sanger sequencing in each iPSC edited single cell clone. For clones with more than one pathogenic *NF2* variant, the cDNA coding sequence of the *NF2* gene was cloned using the Gateway® Gene Cloning System (Invitrogen). The resulting clones were then analysed by Sanger sequencing to determine whether both variants were present on the same allele (cis) or on different alleles (trans).

#### Massive Genotyping

Massive genotyping analysis was assessed using GeneTitan MC Fast Scan Instrument (ThermoFisher Scientific) and NIMBUS Target Preparation Instrument, according to manufacturers’ instructions. The Axiom Precision Medicine Diversity Research Array Plate (PMDA 96-Array Plate) was used to analyze over 830,000 SNPs. All samples were analyzed independently and treated as unpaired samples. The 96-well plate was filled by 92 clinical samples, the PMDA’s positive control and 3 samples from the (+/+), (+/-) and (+/-) cell lines.

#### SNP-array data processing

The CEL files obtained from GeneTItan were all analyzed with Affymetrix Power Tools (APT), following Axiom’s bests practices recommendations for Copy Number Variation (CNV) analysis (https://assets.thermofisher.com/TFS-Assets/GSD/Scientific%C2%A0Guides/axiom-copy-number-data-analysis-guide.pdf).

#### Whole Exome Sequencing

The study of off targets caused by CRISPR Technology was performed by Whole Exome Sequencing (WES) using KAPA HyperExome probes (Roche) and it was sequenced in a NextSeq instrument (Illumina). Analysis of small nucleotide variants (SNVs) was performed using GATK (60). We filtered out those variants that were shared between the (+/-) and (-/-) cells and for those that affect exons in MANE Select transcripts (61).

### IPSCS CHARACTERIZATION

This study was performed at Barcelona Stem Cell Bank as described in Catasús, et al.2025 (45)

#### Study of pluripotency-associated markers

Nuclei were stained with 4’,6-diamidino-2-phenylindole (DAPI). The list of antibodies used is provided in (Tab. S5). Confocal images were acquired using Leica TCS SPE/SP5 microscopes.

#### iPSCs differentiation into the three primary germinal layers through embryoid body (EB)formation

Cells were analyzed by immunofluorescence as described in the *Study of pluripotency-associated markers section*. The list of antibodies used is provided in (Tab. S5). Confocal images were captured using Leica TCS SPE/SP5 microscopes.

#### Alkaline Phosphatase activity

Alkaline Phosphatase Blue Substrate Solution (Sigma) was used to demonstrate iPSC alkaline phosphatase activity.

#### Karyotype determination

Karyotype was evaluated as previously described (62). by G banded metaphase karyotype analysis, at the Hospital San Joan de Deu (Barcelona).

#### iPSCs characterization by Western Blot

Cells were lysed with RIPA buffer (50 mM Tris-HCl (pH 7.4), 150 mM NaCl, 1mM EDTA, 0.5% Igepal CA-630) supplemented with 3mM DTT (Roche), 1mM PMSF (Fluka), 1mM sodium orthovanadate (Sigma), 5mM NaF (Honeywell), 10 ug/ml leupeptin (Sigma), 0.5ug/ml aprotinin (Sigma) and 1xPhosSTOP (Roche). 50 µg of protein were loaded to SDS-PAGE gel (150V) and transferred to PVDF membranes (1 hour 350 mA at 4°C). Odyssey Blocking Buffer TBS (LI-COR) was used to block the membranes. Primary antibodies were incubated overnight at 4°C. Secondary antibodies were posteriorly incubated (1:1000 dilution, LI-COR) for 1h at room temperature and scanned and analyzed using the infrared imaging system LI-COR Odyssey Clx Platform. α-vinculin primary antibody was used to normalize protein expression between samples (Tab. S5).

### NC-SC CHARACTERIZATION

NS-SC characterization were performed as described in Catasús, et al. 2025 (45).

#### Immunofluorescence assay

Cells or spheroids previously fixed, permeabilized and blocked were incubated in primary antibodies (Tab. S5) overnight at 4°C. Secondary antibodies were incubated for 1 hour at RT. Images were acquired using LEICA DMIL6000 microscope and LAS X software.

#### Flow cytometry assay

Collected cells were resuspended in PBS supplemented with 1% BSA and incubated with α-p75 and Hnk 1 primary antibodies were incubated 30 minutes on ice followed by incubation with secondary antibody (Tab. S5). Cells were analyzed using a BD LSR Fortessa SORP flow cytometer, and data was processed with BD FACS Diva 6.2 software.

#### Proliferation assay

After 72 hours, NC cells were treated with 10 µM EdU for 2 hours and subsequently processed using the Click-iT EdU Alexa Fluor 647 Flow Cytometry Assay Kit (Thermo Fisher) according to the manufacturer’s instructions.

#### RNA processing, sequencing and analysis

Total RNA extraction from iPSCs, NC cells and SC-differentiating spheroids was extracted with Maxwell RSC simplyRNA Cells Kit (Promega), following manufacturer’s instructions. RNA was quantified with a Nanodrop 1000 spectrophotometer (Thermo Scientific). The polyA RNA libraries were sequenced on an Illumina Novaseq 6000 in 150 bp pair-end mode. Data was analyzed as described below.

#### Engraftment orthotopic model

2.25·10^6^ NC cells/well were seeded onto AggreWell TM800 24-well plates as described above. After 14 days of differentiation, spheroids were collected and injected into the sciatic nerve of nude mice as described (63). Simultaneously, 1·10^6^ BenMen-1 cells were injected as a control following the same procedure. After 4 months, animals were sacrificed by cervical dislocation, the rear limbs were removed for dissection. The biceps femoris and the gastrocnemius muscles were dissected and fixed in paraffin-embedded for H&E morphological characterization.

### EVALUATION OF THE PMO TREATMENT EFFECT

#### Antisense Phosphorodiamidate Morpholino Oligomers (PMOs) Treatment

PMOs molecules were specifically designed (Tab. S6), synthesized and purified by Gene Tools (Philomath) to induce exon skipping of exon 11. For iPSCs and NC cells, 10^4^ cells/well were seeded in a 12 wells plate. 24h later, medium was replaced with mTSER with 40µM PMOs and 6mM EndoPorter Simple Delivery Reagent (Gene Tools), used as a vehicle. An evaluation of the dose response was performed in previous studies (28). At 72h cells were recollected to extract RNA. When working with SC, 2.25·10^6^ NC cells/well were seeded onto AggreWell TM800 24-well plates as described above. Cells were treated with 2µM of Vivo-Morpholino (Gene Tools) after 7 days of differentiation onto SC. After 3 days of treatment, cells were collected to extract RNA to perform bulk RNAseq and analyze *NF2* levels by RT-PCR.

#### Apoptosis Assay

Apoptosis levels were studied by Annexin V assay. After 7 days of differentiation, cells were treated with Vivo-Morpholino. After 3 days of treatment, 100uL of cell suspension were incubated with Annexin V. The death control was performed by a thermic shock. Images were acquired using LEICA DMIL6000 microscope and LAS X software.

#### Spheroid Size Assay

Spheroid area was measured at 3, 5 and 7 days of PMO treatment. Images were acquired using LEICA DMIL6000 microscope, LAS X software and ImageJ software.

#### Nuclear YAP assay

Spheroids were fixed, permeabilized and blocked. Primary antibody (Tab. S5). was incubated overnight at 4°C. Secondary antibody, was incubated for 1 hour at RT. Nuclei were stained with DAPI (Stem Cell Technologies, 1:1000). Images were acquired using LEICA DMIL6000 microscope and LAS X software. Cytofluorogram was assessed using BIOP JaCoP pluging in ImageJ.

#### Western Blot

After 7 days of differentiation, SC spheroids were treated with PMOs for 3 days and collected for protein extraction. Protein extraction and Western Blot were assessed as described above (Tab. S5). Protein levels of control cells were used to normalize protein expression between samples. Min-mas normalization was applied.

#### RNA processing, sequencing and analysis

Total RNA extraction from untreated SC spheroids, treated with a Control Vivo-Morpholino SC spheroids and treated with specific Vivo-Morpholino SC spheroids during 3, 5 and 7 days, was extracted with Maxwell RSC simplyRNA Cells Kit (Promega), following manufacturer’s instructions. RNA was quantified with a Nanodrop 1000 spectrophotometer (Thermo Scientific). The polyA RNA libraries were sequenced on an Illumina Novaseq 6000 in 150 bp pair-end mode. Sequencing quality was assessed with fastQC and low quality and adapter were trimmed with bbduk (64). STAR (65) was used to map the reads against the human genome version hg38, counts were obtained using featureCounts (66). Raw counts matrices were used for differential expression (DE) analysis by DESeq2 (67), comparisons were made using Wald test. Gene Set Enrichment Analysis (GSEA) has been performed using the fgsea R package (68) ranking values by stat. PCAs were performed on the vst values by DSEQ2 using the top 500 most variable genes. The Single Sample GSEA (ssGSEA) results were obtained using the GSVA R package (69). Gene sets annotation for GSEA and ssGSEA were obtained from MSigDB (70) v2025.1.Hs. All plots derived from the RNA-Seq were done using ggplot2 (71).

#### STRINGdb analysis

We conducted a network association analysis via STRINGdb (72) by checking the association of the off target labelled genes with NF2 in a full STRING network.

#### Functional Enrichment Analysis

Functional enrichment analysis were performed using clusterProfiler (73), for the enrichment, 0.05 and 0.1 as p value and q value cutoffs.

## DATA AVAILABILITY STATEMENT

The authors confirm that the data supporting the findings of this study are available within the article [and/or] its supplementary materials. Raw data for WES and RNAseq is allocated at the European Geno-Phenome Archive (EGA) with accession number XXXXXXXXX and will be made available to external researchers upon reasonable requests.

## ACKNOWLEDGEMENTS

We thank the Spanish Reference Center on Phakomatosis and Hospital Universitari Germans Trias I Pujol’s staff for their collaboration in collecting patient samples, the Hereditary Cancer Group at the IGTP for their help in improving this work, the IGTP Flow Cytometry and Microscopy core facilities, the Mouse Lab at SCT-IDIBELL and the BCN Stem Cell Bank, and their staff for their technical support. We would like to acknowledge the constant support of the different NF lay associations: Asociación de Afectados de Neurofibromatosis, Chromo22, Children’s tumor foundation (CTF) and ACNefi.

## AUTHOR CONTRIBUTIONS

GC: Collection and/or assembly of data; Data analysis and interpretation; Manuscript writing; IJ: Data analysis and interpretation; Manuscript writing; IBo, NC: Collection and/or assembly of data; AQ: In vivo assay data analysis and interpretation; Manuscript writing.. EA: Provision of study material or patients; SP, CL, ES, IBl: Scientific input; EC: Conception and design; Financial support; Manuscript writing; Final approval of manuscript.

