## Supplementary data for "*NF2* lacking exon 11 induced by antisense gene therapy is able to partially recover merlin deficiency on *NF2*-SWN iPSC-derived spheroid model"

#### SUPLEMENTARY INFORMATION

#### Supplementary Material and methods:

##### CELL CULTURE:

*iPSCs culture*
iPSCs were grown on 6-well plates previously coated with growth factor-reduced Matrigel (BD Biosciences) (1:20 dilution) and cultured in mTESR Plus medium (STEMCELL Technologies). When necessary, differentiated cells were removed manually, in brief, a Diascopic Trinocular Microscope was placed into the culture cabin to see the morphology of every cell colony, those colonies presenting differentiated morphology were detached and removed individually. *NF2*(+/-) and *NF2*(+/+) iPSCs lines were split using Accutase (Merk), *NF2*(-/-) cells were split manually, selecting the colonies that presented a pluripotent morphology. All lines were seeded with Rock Inhibitor (STEMCELL Technologies) (1:1000) for 24h. Medium was changed every 24h and maintained at 37ºC and 5% $CO_2$. Cells were frozen with StemMACS Cryo-Brew (Miltenyi Biotec). In this study, the *NF2*(+/+) FiPS line (FiPS Ctrl 1-SV4F-7, (FiPS Ctrl 1-SV4F-7 registered in the Spanish National Stem Cell Bank/ ESiO44C in https://hpscreg.eu/)) was used in all experiments as a control.

*Neural Crest differentiation*
iPSC lines were differentiated to Neural Crest as previously described (1). 9x$10^4$ iPSCs were seeded on matrigel-coated 6-well plates (1:20 dilution) in mTESR Plus medium, 24h later the medium was replaced with Neural Crest Differentiation Media (described below). Medium was changed every day, cells were maintained at 37ºC and 5% CO2. When reaching 80% of confluence, splits were done by detaching the cells enzymatically with Accutase (Merk). After 10 days of differentiation, expression of NC specific markers was studied by flow cytometry and by immunocytochemistry assays. Cells were frozen with FBS + DMSO 10%.

*Neural Crest media*
DMEM:F12 (Gibco) 1:1; 5mg/mL BSA (Sigma); 100U/mL penicillin/ 100mg/mL streptomycin (Gibco); 2mM GlutaMAX (Gibco); 1x MEM non-essential amino acids (Gibco); 1x trace elements A; 1x trace elements B; 1x trace elements C (Corning); 90µM 2-mercaptoethanol (Gibco); 10µg/mL apo-transferrin human(Sigma); 50µg/mL sodium L-ascorbate (Sigma); 10ng/mL Recombinant human Heregulin beta-1 (PeproTech); 200ng/mL Recombinant human IGF-I LR3 (PeproTech); 8ng/mL basic fibroblast growth factor 2 (PeproTech), 2µM CHIR9902 (STEMCELL Technologies) and 20µM SB432542 (STEMCELL Technologies).

*Schwann Cell differentiation in 3D*
NC cells were detached with Accutase, 2.25·$10^6$ cells/well were seeded onto AggreWell TM800 24-well plates (Stem Cell Technologies) in 2mL SCDM (described below) and cultured at 37ºC and 5% CO2. AggreWell TM800 plates were previously treated with Anti-Adherence Rinsing Solution (StemCell technologies). The medium was changed twice a week removing 1 mL and replacing 1 mL of fresh SCDM. On days 7 and 14 spheroids were collected and processed for subsequent analysis. Cells were cultured for a maximum of 14 days.

*Schwann Cell media (SCDM)*
DMEM: F12 (3:1); 100U/ml penicillin/ 100mg/mL streptomycin antibiotics (Gibco); 5µM forskolin (Sigma); 50ng/mL Recombinant human Heregulin beta-1 (Peprotech); 2% N2 supplement (Gibco); 1% FBS (Gibco).

#### IPSC EDITING AND GENOMICS CHARACTERIZATION

*CRISPR/Cas9 gene edition in iPSC lines*

CRISPR/Cas9 editing was conducted using the ArciTect ribonucleoprotein (RNP) system (STEMCELL Technologies). The sgRNA targeted exon 11 of the *NF2* gene (TCGCTCGAGAGAAGCAGATG) and was designed using the Synthego CRISPR Design Tool (CRISPR Design Tool, 2024, v.1.3). Transfection was performed with the TransIT-X2® Dynamic Delivery System (Mirus) following the manufacturer's instructions.

*NF2 gene characterization after CRISPR/Cas gene editing:*

*NF2* exon 11 was screened by Sanger sequencing in each iPSC edited single cell clone. For clones with more than one pathogenic *NF2* variant, the cDNA coding sequence of the *NF2* gene was cloned using the Gateway® Gene Cloning System (Invitrogen). The resulting clones were then analysed by Sanger sequencing to determine whether both variants were present on the same allele (cis) or on different alleles (trans).

*Variant analysis:*

Human Genome Variation Society (www.hgvs.org) nomenclature guidelines were used to name the mutation at the DNA level, its effect at the mRNA level, and the predicted resulting protein. The first nucleotide of the ATG translation initiation codon is denoted position þ1 according to the *NF2* mRNA sequence NM_000268.

*Massive Genotyping*

Massive genotyping analysis was assessed using GeneTitan MC Fast Scan Instrument (ThermoFisher Scientific) and NIMBUS Target Preparation Instrument, according to manufacturers' instructions. The Axiom Precision Medicine Diversity Research Array Plate (PMDA 96-Array Plate) was used to analyze over 830,000 SNPs. All samples were analzed independently and treated as unpaired samples. The 96-well plate was filled by 92 clinical samples, the PMDA's positive control and 3 samples from the (+/+), (+/-) and (+/-) cell lines.

*SNP-array data processing*

The CEL files obtained from GeneTItan were all analyzed with Affymetrix Power Tools (APT), following Axiom's bests practices recommendations for Copy Number Variation (CNV) analysis (https://assets.thermofisher.com/TFS-Assets/GSD/Scientific%C2%A0Guides/axiom-copy-number-data-analysis-guide.pdf). First, all the samples of the plate were filtered by DQC >= 0.82 and call rate >= 97%, no sample on the plate had to be removed after quality assessment which was performed using apt-geno-qc-axiom and apt-genotype-axiom. Samples were genotyped using apt-genotype-axiom and B Allele Frequency (BAF) and Log R Ratio (LRR) were calculated using apt-copynumber-axiom-hmm. The data was denoised and segmented using the DNA copy R package (2) with its default parameters.

*Whole Exome Sequencing*

The study of off targets caused by CRISPR Technology was performed by Whole Exome Sequencing (WES). In brief, KAPA HyperCap technology with KAPA HyperExome Probes (Roche) was used according to manufacturer's instructions and sequenced in a NextSeq instrument (Illumina). Analysis of small nucleotide variants was performed with GATK (3) following the bests practices for germline short variant discovery (https://gatk.broadinstitute.org/hc/en-us/articles/360035535932-Germline-short-variant-discovery-SNPs-Indels). Once the lists of variants were obtained for each cell line, we filtered out those variants that were shared between the (+/-) and (-/-) cells. Variants were filtered for those that affect exons, and which were in MANE Select transcripts (4).

#### IPSCS CHARACTERIZATION

*Study of pluripotency-associated markers*

iPSCs were fixed with 4% paraformaldehyde (PFA), followed by blocking and permeabilization with TBS containing 0.5% Triton X-100 and 6% donkey serum. Primary antibodies were incubated overnight in TBS with 0.1% Triton X-100 and 6% donkey serum. Secondary antibodies were incubated for 2 hours at 37°C. Nuclei were stained with 4',6-diamidino-2-phenylindole (DAPI). The list of antibodies used is provided in (Tab. S5). Confocal images were acquired using Leica TCS SPE/SP5 microscopes. This protocol was performed at Barcelona Stem Cell Bank.

*iPSCs differentiation into the three primary germinal layers through embryoid body (EB)formation*

iPSC colonies were lifted using EDTA and transferred to a 96-well plate with mTeSR-1 medium (Stem Cell Technologies) using a multichannel pipette. The plate was centrifuged at 800g for 10 minutes and incubated at 37°C with 5% $CO_2$ for 24 hours. Early embryoid bodies (EBs) were then transferred to an ultra-low attachment plate in mTeSR-1 medium for additional 24 hours. After this period, EBs were transferred to Matrigel-coated slide flasks and cultured in differentiation medium (described below) for 21-28 days.

- Ectoderm medium: 50% Neurobasal medium, 50% DMEM/F12, 1% N2, 1% B27, 1% Glutamax, and 1% Penicillin-Streptomycin.
- Endoderm medium: Knockout-DMEM, 10% FBS, 1% NEAA, 0.1% β-mercaptoethanol, 1% Glutamax, and 1% Penicillin-Streptomycin (all Gibco).
- Mesoderm medium: Endoderm medium supplemented with 0.5mM ascorbic acid.

Cells were analyzed by immunofluorescence as described in the *Study of pluripotency-associated markers section*. The list of antibodies used is provided in (Tab. S5). Confocal images were captured using Leica TCS SPE/SP5 microscopes. This protocol was performed at Barcelona Stem Cell Bank.

*Alkaline Phosphatase activity*

Alkaline Phosphatase Blue Substrate Solution (Sigma) was used to demonstrate iPSC alkaline phosphatase activity. This protocol was performed at Barcelona Stem Cell Bank.

*Karyotype determination*

iPSC karyotype was assessed by treating cells with colcemid (Gibco) and processed as previously described (5). Karyotype was evaluated by G banded metaphase karyotype analysis, at the Hospital San Joan de Deu (Barcelona).

*Western Blot*

Cells were lysed with RIPA buffer (50 mM Tris-HCl (pH 7.4), 150 mM NaCl, 1mM EDTA, 0.5% Igepal CA-630) supplemented with 3mM DTT (Roche), 1mM PMSF (Fluka), 1mM sodium orthovanadate (Sigma), 5mM NaF (Honeywell), 10 ug/ml leupeptin (Sigma), 0.5ug/ml aprotinin (Sigma) and 1xPhosSTOP (Roche). 50 µg of protein were loaded to SDS-PAGE gel (150V) and transferred to PVDF membranes (1 hour 350 mA at 4°C). Odyssey Blocking Buffer TBS (LI-COR) was used to block the membranes. Primary antibodies were incubated overnight at 4°C. Secondary antibodies were posteriorly incubated (1:1000 dilution, LI-COR) for 1h at room temperature and scanned and analyzed using the infrared imaging system LI-COR Odyssey Clx Platform. α-vinculin primary antibody was used to normalize protein expression between samples (Tab. S5).

#### NC-SC CHARACTERIZATION

*Immunofluorescence assay*

Cells or spheroids were fixed in 4% paraformaldehyde in PBS for 15 minutes at room temperature (RT), followed by permeabilization with 0.1% Triton-X 100 in PBS for 10 minutes at RT. Blocking was carried out with 10% FBS in PBS for 15 minutes at RT. Primary antibodies (Tab. S5). were incubated overnight at 4°C. Secondary antibodies, Alexa Fluor 488 and Alexa Fluor 568, were incubated for 1 hour at RT. Nuclei were stained with DAPI (Stem Cell Technologies, 1:1000). Images were acquired using LEICA DMIL6000 microscope and LAS X software.

*Flow cytometry assay*

$1.10^6$ cells, previously detached using Accutase, were resuspended in PBS supplemented with 1% BSA and incubated with α-p75 primary antibody, followed by incubation with Alexa Fluor 568 conjugated secondary antibody. Subsequently, the cells were incubated with α-Hnk1 primary antibody, followed by Alexa Fluor 488 conjugated secondary antibody (Tab. S5). All antibodies were incubated 30 minutes on ice. Cells were analyzed using a BD LSR Fortessa SORP flow cytometer, and data was processed with BD FACS Diva 6.2 software.

*Proliferation assay*

$3.10^5$ cells were seeded onto 6-well plates pre-coated with Matrigel and cultured with Neural Crest Differentiation Media. After 72 hours, cells were treated with 10 µM EdU for 2 hours and subsequently processed using the Click-iT EdU Alexa Fluor 647 Flow Cytometry Assay Kit (Thermo Fisher) according to the manufacturer's instructions. Cells were also stained with DAPI to assess DNA content. Data acquisition and analysis were performed using a BD LSR Fortessa SORP flow cytometer, results were processed using BD FACSDiva 6.2 software.

*Scratch Assay*

Cells were seeded onto 6-well plates coated with Matrigel. Once they reached approximately 80% confluence, a scratch was made using a pipette tip to create a gap. Cell migration was assessed by capturing images of the same region at 6 and 24 hours post-scratch. Images were acquired using a LEICA DMIL6000 microscope and analysed with LAS X software.

*RNA processing, sequencing and analysis*

Total RNA extraction from iPSCs, NC cells and SC-differentiating spheroids was extracted with Maxwell RSC simplyRNA Cells Kit (Promega), following manufacturer's instructions. RNA was quantified with a Nanodrop 1000 spectrophotometer (Thermo Scientific). The polyA RNA libraries were sequenced on an Illumina Novaseq 6000 in 150 bp pair-end mode. Data was analyzed as described below.

*Engraftment orthotopic model*

$2.25 \cdot 10^6$ NC cells/well were seeded onto AggreWell TM800 24-well plates as described above. After 14 days of differentiation, spheroids were collected and injected into the sciatic nerve of nude mice as described (6). Simultaneously, $1 \cdot 10^6$ BenMen-1 cells were injected as a control following the same procedure. After 4 months, animals were sacrificed by cervical dislocation, the rear limbs were removed for dissection. The biceps femoris and the gastrocnemius muscles were dissected and fixed in paraffin-embedded for H&E morphological characterization.

#### EVALUATION OF THE PMO TREATMENT EFFECT

*Antisense Phosphorodiamidate Morpholino Oligomers Treatment*

Antisense Phosphorodiamidate Morpholino Oligomer (PMOs) molecules were specifically designed (Tab. S6), synthesized and purified by Gene Tools (Philomath) to induce exon skipping of exon 11. For iPSCs and NC cells, $10^4$ cells/well were seeded in a 12 wells plate. 24h later, medium was replaced with mTSER with 40µM PMOs and 6mM EndoPorter Simple Delivery Reagent (Gene Tools), used as a vehicle. An evaluation of the dose response was performed in previous studies (7). At 72h cells were recollected to extract RNA. When working with SC, $2.25 \cdot 10^6$ NC cells/well were seeded onto AggreWell TM800 24-well plates as described above. Cells were treated with 2µM of Vivo-Morpholino (Gene Tools) after 7 days of differentiation onto SC. After 3 days of treatment, cells were collected to extract RNA.

*RT-PCR*

RNA was extracted with Maxwell RSC simplyRNA cells kit, from Maxwell technologies (Promega). Afterwards, total RNA was quantified using Nanodrop (Thermofisher) and 0.5µg were reverse transcribed by RT-PCR, performed using SuperScript III reverse transcriptase (Invitrogen) and random hexamers (Invitrogen) to obtain cDNA. Subsequently, a PCR of the obtained cDNA was performed targeting a fragment that included from exon 9 to 12. PCR conditions were 35 cycles with an annealing temperature of 59ºC. Samples were posteriorly Sanger sequenced by Macrogen.

*Apoptosis Assay*

Apoptosis levels were studied by Annexin V assay. $2.25 \cdot 10^6$ NC cells/well were seeded onto AggreWell TM800 24-well plates as described above. After 7 days of differentiation, cells were treated with a dose of 2µM Vivo-Morpholino. After 3 days of treatment, spheroids were collected and resuspended with Annexin V Binding Buffer. 100uL of cell suspension were incubated with 5uL of Annexin V 15min at RT. Nuclei was stained with DAPI (Stem Cell Technologies, 1:1000). The death control was performed by a thermic shock of 20min at 70ºC and 15min at 4ºC. Images were acquired using LEICA DMIL6000 microscope and LAS X software.

*Spheroid Size Assay*

Spheroids were generated by seeding $2.25 \cdot 10^6$ NC cells/well onto AggreWell TM800 24-well plates as described above. After 7 days of differentiation, spheroids were treated with 2µM of Vivo-Morpholino. Spheroid area was measured at 3, 5 and 7 days of treatment. Images were acquired using LEICA DMIL6000 microscope, LAS X software and ImageJ software.

*Nuclear YAP assay*

Spheroids were fixed in 4% paraformaldehyde in PBS for 15 minutes at room temperature (RT), followed by permeabilization with 0.1% Triton-X 100 in PBS for 10 minutes at RT. Blocking was carried out with 10% FBS in PBS for 15 minutes at RT. Primary antibody (Tab.... S5). was incubated overnight at 4°C. Secondary antibody, Alexa Fluor 488, was incubated for 1 hour at RT. Nuclei were stained with DAPI (Stem Cell Technologies, 1:1000). Images were acquired using LEICA DMIL6000 microscope and LAS X software. Nuclear YAP1 was studied by evaluating the co-localization of YAP1 and DAPI stainings. Cytofluorogram was assessed using BIOP JaCoP pluging in ImageJ, auto-thresholds used were Default (DAPI) and Moments (YAP1).

*Western Blot*

After 7 days of differentiation, SC spheroids were treated with 2uM for 3 days and collected for protein extraction. Protein extraction and Western Blot were assessed as described above (Tab. S5). Protein levels of control cells were used to normalize protein expression between samples. Min-mas normalization was applied.

*RNA processing, sequencing and analysis*

Total RNA extraction from untreated SC spheroids, treated with a Control Vivo-Morpholino SC spheroids and treated with specific Vivo-Morpholino SC spheroids during 3, 5 and 7 days, was extracted with Maxwell RSC simplyRNA Cells Kit (Promega), following manufacturer's instructions. RNA was quantified with a Nanodrop 1000 spectrophotometer (Thermo Scientific). The polyA RNA libraries were sequenced on an Illumina Novaseq 6000 in 150 bp pair-end mode. Reads' quality was assessed by using fastQC (Andrews, S., (2010)). Reads were trimmed with the bbduk program (8) using the following parameters (ktrim=r k=23 mink=11 hdist=1 tpe tbo qtrim=rl trimq=20 minlen=23). Quality and adapter trimmed reads were aligned with STAR (9) against the human genome primary assembly (version hg38) which was obtained via Ensembl (https://ftp.ensembl.org/pub/release-114/fasta/homo_sapiens/dna/). Alignment was performed using the STAR's default parameters. Matrix counts were obtained using featureCounts (10) with the –-primary –p –s 1 –C –B flags. Raw counts matrices were used for differential expression (DE) analysis by DESeq2 (11). Comparisons were made using the Wald significance test and p values adjustment (FDR) was performed with the Benjamini-Hochberg correction. To classify a gene as differentially expressed, we considered those with an adjusted p value lower than 0.01 and with an absolute value of LogFC greater than 1. For the modelling of the data in DESeq2, we used the following model for the differentiating cells was: *~genotype + celltype + genotype:celltype* and for the treated cells experiment it was: *~treatment + days + genotype + genotype:treatment + treatment:days.* Gene Set Enrichment Analysis (GSEA) analysis has been performed using the fgsea R package (12), genes were ranked based on the *stat* value of the DESeq2 results, filtering out all the genes with no stat value available, results were filtered by FDR 0.05. PCA were performed on the vst values given by DESeq2, the top 500 genes at the SD level for the analysis. The Single Sample GSEA (ssGSEA) results were obtained using the GSVA R package (13). Gene sets annotation for GSEA and ssGSEA were obtained from MSigDB (14) v2025.1.Hs. All plots derived from the RNA-Seq were done using ggplot2 (15).

*STRINGdb analysis*

The relation of off target labelled genes with NF2 was investigated, to do so we conducted an association functional analysis on STRINGdb (16). The network type was set to be a full STRING network, Interactions were set to have a minimum confidence of 0.7 and the enables sources of interaction were: Text mining, Experiments, Database and Co-expression.

*Functional Enrichment Analysis*

To conduct functional enrichment analysis, we used the cluster Profiler R (17) package, p value and q value cutoffs were set as 0.05 and 0.1 respectively and only pathways with a minimum of 10 present genes were considered for the analysis. Only protein coding genes were considered for the enrichment analysis that was conducted with the whole proteome as background.

#### Supplementary tables

**Supplementary table S1.** CRISPR-generated lines characterization

| Clone Name | *NF2* genotype | Genomic (g.) | Coding (c.) | Protein (p.) | Affected exon |
|---|---|---|---|---|---|
| 2H9 | *NF2* (+/-) | g.73302_73305delAGAT | c.1031_1034delAGAT | p.(Gln344Argfs*19) | 11 |
| | | WT | WT | WT | |
| 2F6 | *NF2* (-/-) | g.73303delG | c.1032delG | p.(Met345*) | 11 |
| | | g.73303dupG | c.1032dupG | p.(Met345Aspfs*6) | |

**Supplementary Table S2.** WES analysis of the CRISPR-generated lines

| ***NF2* (+/-) [2H9]** | | | | |
|---|---|---|---|---|
| **Gene Symbol** | **Variant Classification** | **Variant Type** | **Genome Change** | **Observations** |
| **ANKRD30BP2** | intronic | INS | g.chr21:13065558_13065559insT | Not related to disease |
| **ZNG1C** | frameshift | DEL | g.chr9:68285995_68285996insA | Not related to disease |
| **MKI67P1** | intronic | SNP | g.chrX:46560997delT | Not related to disease |
| **R3HDM2P2** | intronic | SNP | g.chr6:104019748C>G | Not related to disease |
| **CYCSP20** | intronic | SNP | g.chr7:129117515T>A | Not related to disease |
| **TRBV11-2** | intronic | SNP | g.chr7:142433897T>C | Not related to disease |
| **CARS1P1** | intronic | SNP | g.chr15:68750234G>A | Not related to disease |
| **ANAPC1** | nonsense | SNP | g.chr2:111856852G>A | Related to Rothmund-Thomson Syndrome type 1 (TRS type 1), recessive disease. |
| **GYPE** | missense | SNP | g.chr4:143880509C>T | Not related to disease |
| **EMB** | de novo start in frame | SNP | g.chr5:50441293A>C | Not related to disease |
| **UNKL** | frameshift | INS | g.chr16:1397277_1397278ins | Not related to disease |
| **UNKL** | frameshift | INS | g.chr16:1397281_1397282ins | Not related to disease |
| **ENSG00000227291** | intronic | DEL | g.chr2:117181177delT | Not related to disease |
| **SDCBPP2** | intronic | INS | g.chr8:69944278_69944279insT | Not related to disease |
| **PRDX2P2** | intronic | INS | g.chr10:34965748_34965749insT | Not related to disease |
| **NKRF** | frameshift | INS | g.chrX:119605952_119605953insG | Not related to disease |
| **HSPD1P12** | intronic | INS | g.chr12:8016406_8016407insT | Not related to disease |
| **ENSG00000258639** | intronic | SNP | g.chr14:48045482G>C | Not related to disease |

| *NF2* (-/-) [2F6] | | | | |
|---|---|---|---|---|
| **Gene Symbol** | **Variant Classification** | **Variant Type** | **Genome Change** | **Observations** |
| **ANKRD30BP2** | intronic | INS | g.chr21:13065558_13065559insT | Not related to disease |
| **ZNG1C** | frameshift | INS | g.chr9:68285995_68285996insA | Not related to disease |
| **MKI67P1** | intronic | DEL | g.chrX:46560997delT | Not related to disease |
| **R3HDM2P2** | intronic | SNP | g.chr6:104019748C>G | Not related to disease |
| **CYCSP20** | intronic | SNP | g.chr7:129117515T>A | Not related to disease |
| **TRBV11-2** | intronic | SNP | g.chr7:142433897T>C | Not related to disease |
| **CARS1P1** | intronic | SNP | g.chr15:68750234G>A | Not related to disease |
| **ANAPC1** | nonsense | SNP | g.chr2:111856852G>A | Related to Rothmund-Thomson Syndrome type 1 (TRS type 1), recessive disease. |
| **GYPE** | missense | SNP | g.chr4:143880509C>T | Not related to disease |
| **EMB** | de novo start in frame | SNP | g.chr5:50441293A>C | Not related to disease |
| **RPS29P27** | intronic | INS | g.chr19:39697905_39697906insG | Not related to disease |
| **LRRC37A14P** | intronic | INS | g.chr22:41190188_41190189insAA | Not related to disease |
| **ENSG00000250609** | intronic | DEL | g.chr4:154523675delT | Not related to disease |
| **SEPTIN14P1** | intronic | DEL | g.chr7:63148831delG | Not related to disease |
| **ANKRD18CP** | intronic | SNP | g.chr9:97179996C>G | Not related to disease |
| **GXYLT1** | nonsense | SNP | g.chr12:42144550C>A | Not related to disease |
| **ENSG00000257398** | intronic | SNP | g.chr12:107845965T>A | Not related to disease |
| **ENSG00000227098** | intronic | SNP | g.chr2:176843497A>G | Not related to disease |

**Supplementary table S3.** Orthotopic model experimental design.

| Experiment | Cell line | Genotype | Mouse ID | Sex |
|---|---|---|---|---|
| E95/24 | FiPs | *NF2* (+/+) | B1R (R) | Male |
| E95/24 | FiPs | *NF2* (+/+) | B1L (R) | Male |
| E95/24 | FiPs | *NF2* (+/+) | D1R (R) | Female |
| E95/24 | BenMen- 1 | *NF2* (-/-) | D1L (R) | Female |
| E95/24 | BenMen- 1 | *NF2* (-/-) | DLR (L) | Female |
| E95/24 | BenMen- 1 | *NF2* (-/-) | DLR (R) | Female |
| E95/24 | 2H9 | *NF2* (+/-) | B1R (L) | Male |
| E95/24 | 2H9 | *NF2* (+/-) | B1L (L) | Male |
| E95/24 | 2H9 | *NF2* (+/-) | BRL (L) | Male |
| E95/24 | 2F6 | *NF2* (-/-) | C1L (L) | Female |
| E95/24 | 2F6 | *NF2* (-/-) | ARL (L) | Male |
| E95/24 | 2F6 | *NF2* (-/-) | A1R (L) | Male |

**Supplementary table S4.** Analysis of unspecific effect of PMO treatment.

| *NF2* (+/-) [2H9] | | | *NF2* (-/-) [2F6] | | |
|---|---|---|---|---|---|
| **gene_id** | **HGNC_symbol** | **Biotype** | **gene_id** | **HGNC_symbol** | **Biotype** |
| ENSG00000198695 | MT-ND6 | protein_coding | ENSG00000198695 | MT-ND6 | protein_coding |
| ENSG00000211459 | MT-RNR1 | Mt_rRNA | ENSG00000211459 | MT-RNR1 | Mt_rRNA |
| ENSG00000249119 | MTND6P4 | processed_pseudogene | ENSG00000249119 | MTND6P4 | processed_pseudogene |
| ENSG00000100292 | HMOX1 | protein_coding | ENSG00000210196 | MT-TP | Mt_tRNA |
| ENSG00000210049 | MT-TF | Mt_tRNA | ENSG00000270230 | MTND6P22 | processed_pseudogene |
| ENSG00000132274 | TRIM22 | protein_coding | ENSG00000210100 | MT-TI | Mt_tRNA |
| ENSG00000136048 | DRAM1 | protein_coding | ENSG00000210151 | MT-TS1 | Mt_tRNA |
| ENSG00000106991 | ENG | protein_coding | ENSG00000290862 | lncRNA | |
| ENSG00000257335 | MGAM | protein_coding | ENSG00000173805 | HAP1 | protein_coding |
| ENSG00000289474 | | lncRNA | ENSG00000214900 | LINC01588 | lncRNA |
| ENSG00000118515 | SGK1 | protein_coding | ENSG00000172183 | ISG20 | protein_coding |
| ENSG00000026103 | FAS | protein_coding | ENSG00000146216 | TTBK1 | protein_coding |
| ENSG00000210100 | MT-TI | Mt_tRNA | ENSG00000161011 | SQSTM1 | protein_coding |
| ENSG00000290862 | | lncRNA | ENSG00000140961 | OSGIN1 | protein_coding |
| ENSG00000067066 | SP100 | protein_coding | ENSG00000134955 | SLC37A2 | protein_coding |
| ENSG00000101665 | SMAD7 | protein_coding | ENSG00000221955 | SLC12A8 | protein_coding |
| ENSG00000151012 | SLC7A11 | protein_coding | ENSG00000276168 | RN7SL1 | misc_RNA |
| ENSG00000294499 | | lncRNA | ENSG00000162772 | ATF3 | protein_coding |
| ENSG00000180353 | HCLS1 | protein_coding | ENSG00000175505 | CLCF1 | protein_coding |
| ENSG00000267898 | | lncRNA | ENSG00000143507 | DUSP10 | protein_coding |
| ENSG00000135919 | SERPINE2 | protein_coding | ENSG00000129990 | SYT5 | protein_coding |
| ENSG00000140044 | JDP2 | protein_coding | ENSG00000144455 | SUMF1 | protein_coding |
| ENSG00000136531 | SCN2A | protein_coding | ENSG00000100439 | ABHD4 | protein_coding |

| | | | | | |
|---|---|---|---|---|---|
| ENSG00000164220 | F2RL2 | protein_coding | ENSG00000004846 | ABCB5 | protein_coding |
| ENSG00000140961 | OSGIN1 | protein_coding | ENSG00000105711 | SCN1B | protein_coding |
| ENSG00000135299 | ANKRD6 | protein_coding | ENSG00000101665 | SMAD7 | protein_coding |
| ENSG00000170323 | FABP4 | protein_coding | ENSG00000197405 | C5AR1 | protein_coding |
| ENSG00000204421 | LY6G6C | protein_coding | ENSG00000099822 | HCN2 | protein_coding |
| ENSG00000205592 | MUC19 | protein_coding | ENSG00000203805 | PLPP4 | protein_coding |
| ENSG00000113749 | HRH2 | protein_coding | ENSG00000258479 | LINC00640 | lncRNA |
| ENSG00000138670 | RASGEF1B | protein_coding | ENSG00000124299 | PEPD | protein_coding |
| ENSG00000106366 | SERPINE1 | protein_coding | ENSG00000185022 | MAFF | protein_coding |
| ENSG00000006128 | TAC1 | protein_coding | ENSG00000167772 | ANGPTL4 | protein_coding |
| ENSG00000099822 | HCN2 | protein_coding | ENSG00000245532 | NEAT1 | lncRNA |
| ENSG00000116285 | ERRFI1 | protein_coding | ENSG00000185896 | LAMP1 | protein_coding |
| ENSG00000117226 | GBP3 | protein_coding | ENSG00000121577 | POPDC2 | protein_coding |
| ENSG00000304089 | | lncRNA | ENSG00000157227 | MMP14 | protein_coding |
| ENSG00000173805 | HAP1 | protein_coding | ENSG00000150630 | VEGFC | protein_coding |
| ENSG00000198108 | CHSY3 | protein_coding | ENSG00000143641 | GALNT2 | protein_coding |
| ENSG00000174749 | FAM241A | protein_coding | ENSG00000160211 | G6PD | protein_coding |
| ENSG00000198774 | RASSF9 | protein_coding | ENSG00000168621 | GDNF | protein_coding |
| ENSG00000105538 | RASIP1 | protein_coding | ENSG00000125730 | C3 | protein_coding |
| ENSG00000118946 | PCDH17 | protein_coding | ENSG00000099860 | GADD45B | protein_coding |
| ENSG00000150630 | VEGFC | protein_coding | ENSG00000169282 | KCNAB1 | protein_coding |
| ENSG00000197405 | C5AR1 | protein_coding | ENSG00000138166 | DUSP5 | protein_coding |
| ENSG00000136205 | TNS3 | protein_coding | ENSG00000142227 | EMP3 | protein_coding |
| ENSG00000242732 | RTL5 | protein_coding | ENSG00000106991 | ENG | protein_coding |
| ENSG00000118985 | ELL2 | protein_coding | ENSG00000205336 | ADGRG1 | protein_coding |
| ENSG00000166342 | NETO1 | protein_coding | ENSG00000100060 | MFNG | protein_coding |
| ENSG00000116016 | EPAS1 | protein_coding | ENSG00000067182 | TNFRSF1A | protein_coding |

| | | | | | |
|---|---|---|---|---|---|
| ENSG00000103175 | WFDC1 | protein_coding | ENSG00000185825 | BCAP31 | protein_coding |
| ENSG00000196611 | MMP1 | protein_coding | ENSG00000092068 | SLC7A8 | protein_coding |
| ENSG00000287763 | | lncRNA | ENSG00000135931 | ARMC9 | protein_coding |
| ENSG00000161011 | SQSTM1 | protein_coding | ENSG00000105880 | DLX5 | protein_coding |
| ENSG00000176046 | NUPR1 | protein_coding | ENSG00000137880 | GCHFR | protein_coding |
| ENSG00000134259 | NGF | protein_coding | ENSG00000122012 | SV2C | protein_coding |
| ENSG00000152049 | KCNE4 | protein_coding | ENSG00000136048 | DRAM1 | protein_coding |
| ENSG00000130513 | GDF15 | protein_coding | ENSG00000287078 | lncRNA | |
| ENSG00000120149 | MSX2 | protein_coding | ENSG00000174939 | ASPHD1 | protein_coding |
| ENSG00000100439 | ABHD4 | protein_coding | ENSG00000126368 | NR1D1 | protein_coding |
| ENSG00000196954 | CASP4 | protein_coding | ENSG00000100985 | MMP9 | protein_coding |
| ENSG00000105329 | TGFB1 | protein_coding | ENSG00000178233 | TMEM151B | protein_coding |
| ENSG00000273415 | LINC02725 | lncRNA | ENSG00000132274 | TRIM22 | protein_coding |
| ENSG00000183379 | SYNDIG1L | protein_coding | ENSG00000142669 | SH3BGRL3 | protein_coding |
| ENSG00000050730 | TNIP3 | protein_coding | ENSG00000138311 | ZNF365 | protein_coding |
| ENSG00000204516 | MICB | protein_coding | ENSG00000006016 | CRLF1 | protein_coding |
| ENSG00000244694 | PTCHD4 | protein_coding | ENSG00000118985 | ELL2 | protein_coding |
| ENSG00000130522 | JUND | protein_coding | ENSG00000213694 | S1PR3 | protein_coding |
| ENSG00000125730 | C3 | protein_coding | ENSG00000162734 | PEA15 | protein_coding |
| ENSG00000183323 | CCDC125 | protein_coding | ENSG00000171435 | KSR2 | protein_coding |
| ENSG00000286177 | | lncRNA | ENSG00000108828 | VAT1 | protein_coding |
| ENSG00000212722 | KRTAP4-9 | protein_coding | ENSG00000011422 | PLAUR | protein_coding |
| ENSG00000092068 | SLC7A8 | protein_coding | ENSG00000095752 | IL11 | protein_coding |
| ENSG00000111981 | ULBP1 | protein_coding | ENSG00000054356 | PTPRN | protein_coding |
| ENSG00000167772 | ANGPTL4 | protein_coding | ENSG00000150551 | LYPD1 | protein_coding |
| ENSG00000166923 | GREM1 | protein_coding | ENSG00000163191 | S100A11 | protein_coding |
| ENSG00000163430 | FSTL1 | protein_coding | ENSG00000128564 | VGF | protein_coding |

| | | | | | |
|---|---|---|---|---|---|
| ENSG00000104435 | STMN2 | protein_coding | ENSG00000107731 | UNC5B | protein_coding |
| ENSG00000150938 | CRIM1 | protein_coding | ENSG00000105329 | TGFB1 | protein_coding |
| ENSG00000287828 | | lncRNA | ENSG00000070404 | FSTL3 | protein_coding |
| ENSG00000163132 | MSX1 | protein_coding | ENSG00000071553 | ATP6AP1 | protein_coding |
| ENSG00000204264 | PSMB8 | protein_coding | ENSG00000116299 | ELAPOR1 | protein_coding |
| ENSG00000138135 | CH25H | protein_coding | ENSG00000164733 | CTSB | protein_coding |
| ENSG00000059728 | MXD1 | protein_coding | ENSG00000143322 | ABL2 | protein_coding |
| ENSG00000196104 | SPOCK3 | protein_coding | ENSG00000164949 | GEM | protein_coding |
| ENSG00000132170 | PPARG | protein_coding | ENSG00000182492 | BGN | protein_coding |
| ENSG00000102034 | ELF4 | protein_coding | ENSG00000223874 | TESHL | lncRNA |
| ENSG00000135318 | NT5E | protein_coding | ENSG00000065054 | NHERF2 | protein_coding |
| ENSG00000289961 | | lncRNA | ENSG00000273415 | LINC02725 | lncRNA |
| ENSG00000286162 | | lncRNA | ENSG00000134072 | CAMK1 | protein_coding |
| ENSG00000164251 | F2RL1 | protein_coding | ENSG00000131188 | PRR7 | protein_coding |
| ENSG00000206337 | HCP5 | lncRNA | ENSG00000100228 | RAB36 | protein_coding |
| ENSG00000251301 | LINC02384 | lncRNA | ENSG00000305609 | lncRNA | |
| ENSG00000163624 | CDS1 | protein_coding | ENSG00000115756 | HPCAL1 | protein_coding |
| ENSG00000171451 | DSEL | protein_coding | ENSG00000118402 | ELOVL4 | protein_coding |
| ENSG00000182492 | BGN | protein_coding | ENSG00000128016 | ZFP36 | protein_coding |
| ENSG00000184584 | STING1 | protein_coding | ENSG00000167601 | AXL | protein_coding |
| ENSG00000143127 | ITGA10 | protein_coding | ENSG00000131711 | MAP1B | protein_coding |
| ENSG00000060140 | STYK1 | protein_coding | ENSG00000109089 | CDR2L | protein_coding |
| ENSG00000198829 | SUCNR1 | protein_coding | ENSG00000289029 | lncRNA | |
| ENSG00000144891 | AGTR1 | protein_coding | ENSG00000132170 | PPARG | protein_coding |
| ENSG00000288765 | | lncRNA | ENSG00000087494 | PTHLH | protein_coding |
| ENSG00000171621 | SPSB1 | protein_coding | ENSG00000104267 | CA2 | protein_coding |
| ENSG00000157227 | MMP14 | protein_coding | ENSG00000062716 | VMP1 | protein_coding |

| | | | | | |
|---|---|---|---|---|---|
| ENSG00000300781 | | lncRNA | ENSG00000102265 | TIMP1 | protein_coding |
| ENSG00000245532 | NEAT1 | lncRNA | ENSG00000167173 | C15orf39 | protein_coding |
| ENSG00000138311 | ZNF365 | protein_coding | ENSG00000120471 | TP53AIP1 | protein_coding |
| ENSG00000164035 | EMCN | protein_coding | ENSG00000105825 | TFPI2 | protein_coding |
| ENSG00000291089 | | lncRNA | ENSG00000136856 | SLC2A8 | protein_coding |
| ENSG00000131015 | ULBP2 | protein_coding | ENSG00000100065 | CARD10 | protein_coding |
| ENSG00000181195 | PENK | protein_coding | ENSG00000179954 | SSC5D | protein_coding |
| ENSG00000125430 | HS3ST3B1 | protein_coding | ENSG00000242802 | AP5Z1 | protein_coding |
| ENSG00000237172 | B3GNT9 | protein_coding | ENSG00000186111 | PIP5K1C | protein_coding |
| ENSG00000041353 | RAB27B | protein_coding | ENSG00000177301 | KCNA2 | protein_coding |
| ENSG00000168938 | PPIC | protein_coding | ENSG00000132692 | BCAN | protein_coding |
| ENSG00000095739 | BAMBI | protein_coding | ENSG00000048540 | LMO3 | protein_coding |
| ENSG00000270069 | MIR222HG | lncRNA | ENSG00000163873 | GRIK3 | protein_coding |
| ENSG00000147655 | RSPO2 | protein_coding | ENSG00000144677 | CTDSPL | protein_coding |
| ENSG00000278464 | | lncRNA | ENSG00000261728 | lncRNA | |
| ENSG00000250337 | PURPL | lncRNA | ENSG00000154274 | C4orf19 | protein_coding |
| ENSG00000101577 | LPIN2 | protein_coding | ENSG00000271327 | lncRNA | |
| ENSG00000122986 | HVCN1 | protein_coding | ENSG00000183508 | TENT5C | protein_coding |
| ENSG00000196932 | TMEM26 | protein_coding | ENSG00000124374 | PAIP2B | protein_coding |
| ENSG00000251254 | GTF2F2P1 | processed_pseudogene | ENSG00000150275 | PCDH15 | protein_coding |
| ENSG00000064012 | CASP8 | protein_coding | ENSG00000177807 | KCNJ10 | protein_coding |
| ENSG00000019549 | SNAI2 | protein_coding | ENSG00000112280 | COL9A1 | protein_coding |
| ENSG00000182253 | SYNM | protein_coding | ENSG00000162654 | GBP4 | protein_coding |
| ENSG00000280219 | | TEC | ENSG00000294479 | lncRNA | |
| ENSG00000143590 | EFNA3 | protein_coding | ENSG00000147145 | LPAR4 | protein_coding |
| ENSG00000166289 | PLEKHF1 | protein_coding | ENSG00000115474 | KCNJ13 | protein_coding |
| ENSG00000198003 | ODAD3 | protein_coding | ENSG00000068650 | ATP11A | protein_coding |

| | | | | | |
|---|---|---|---|---|---|
| ENSG00000168779 | SHOX2 | protein_coding | ENSG00000171843 | MLLT3 | protein_coding |
| ENSG00000139351 | SYCP3 | protein_coding | ENSG00000231252 | lncRNA | |
| ENSG00000279086 | | TEC | ENSG00000160862 | AZGP1 | protein_coding |
| ENSG00000184226 | PCDH9 | protein_coding | ENSG00000077616 | NAALAD2 | protein_coding |
| ENSG00000170382 | LRRN2 | protein_coding | ENSG00000088970 | KIZ | protein_coding |
| ENSG00000176406 | RIMS2 | protein_coding | ENSG00000188580 | NKAIN2 | protein_coding |
| ENSG00000253161 | LINC01605 | lncRNA | ENSG00000226476 | LINC01748 | lncRNA |
| ENSG00000135312 | HTR1B | protein_coding | ENSG00000157734 | SNX22 | protein_coding |
| ENSG00000107485 | GATA3 | protein_coding | ENSG00000286608 | lncRNA | |
| ENSG00000111859 | NEDD9 | protein_coding | ENSG00000250305 | TRMT9B | protein_coding |
| ENSG00000170561 | IRX2 | protein_coding | ENSG00000100784 | RPS6KA5 | protein_coding |
| ENSG00000229931 | ATXN1-AS1 | lncRNA | ENSG00000127084 | FGD3 | protein_coding |
| ENSG00000151617 | EDNRA | protein_coding | ENSG00000272870 | SAP30-DT | lncRNA |
| ENSG00000164949 | GEM | protein_coding | ENSG00000297519 | lncRNA | |
| ENSG00000164588 | HCN1 | protein_coding | ENSG00000106069 | CHN2 | protein_coding |
| ENSG00000305511 | | lncRNA | ENSG00000247809 | NR2F2-AS1 | lncRNA |
| ENSG00000102290 | PCDH11X | protein_coding | ENSG00000264449 | lncRNA | |
| ENSG00000168621 | GDNF | protein_coding | ENSG00000280145 | lncRNA | |
| ENSG00000150977 | RILPL2 | protein_coding | ENSG00000162669 | HFM1 | protein_coding |
| ENSG00000166670 | MMP10 | protein_coding | ENSG00000270207 | lncRNA | |
| ENSG00000274010 | ZFP91P1 | processed_pseudogene | ENSG00000136098 | NEK3 | protein_coding |
| ENSG00000128016 | ZFP36 | protein_coding | ENSG00000283897 | lncRNA | |
| ENSG00000107562 | CXCL12 | protein_coding | ENSG00000139354 | GAS2L3 | protein_coding |
| ENSG00000228526 | MIR34AHG | lncRNA | ENSG00000114805 | PLCH1 | protein_coding |
| ENSG00000107984 | DKK1 | protein_coding | ENSG00000196951 | SCOC-AS1 | lncRNA |
| ENSG00000279041 | | TEC | ENSG00000284624 | lncRNA | |
| ENSG00000289201 | | lncRNA | ENSG00000139132 | FGD4 | protein_coding |

| | | | | | |
|---|---|---|---|---|---|
| ENSG00000136052 | SLC41A2 | protein_coding | ENSG00000165868 | HSPA12A | protein_coding |
| ENSG00000112175 | BMP5 | protein_coding | ENSG00000117115 | PADI2 | protein_coding |
| ENSG00000128203 | ASPHD2 | protein_coding | ENSG00000249577 | processed_pseudogene | |
| ENSG00000283183 | | lncRNA | ENSG00000251034 | lncRNA | |
| ENSG00000290879 | | lncRNA | ENSG00000196482 | ESRRG | protein_coding |
| ENSG00000128422 | KRT17 | protein_coding | ENSG00000291187 | REXO6P | lncRNA |
| ENSG00000165507 | DEPP1 | protein_coding | ENSG00000018408 | WWTR1 | protein_coding |
| ENSG00000181291 | TMEM132E | protein_coding | ENSG00000283638 | MIR106AHG | lncRNA |
| ENSG00000266916 | ZNF793-AS1 | lncRNA | ENSG00000277135 | lncRNA | |
| ENSG00000117318 | ID3 | protein_coding | ENSG00000272983 | lncRNA | |
| ENSG00000138061 | CYP1B1 | protein_coding | ENSG00000188783 | PRELP | protein_coding |
| ENSG00000162631 | NTNG1 | protein_coding | ENSG00000140876 | NUDT7 | protein_coding |
| ENSG00000150551 | LYPD1 | protein_coding | ENSG00000035499 | DEPDC1B | protein_coding |
| ENSG00000168140 | VASN | protein_coding | ENSG00000299851 | lncRNA | |
| ENSG00000164749 | HNF4G | protein_coding | ENSG00000163558 | PRKCI | protein_coding |
| ENSG00000145990 | GFOD1 | protein_coding | ENSG00000150625 | GPM6A | protein_coding |
| ENSG00000186377 | CYP4X1 | protein_coding | ENSG00000214870 | LINC02981 | lncRNA |
| ENSG00000187135 | VSTM2B | protein_coding | ENSG00000108984 | MAP2K6 | protein_coding |
| ENSG00000168016 | TRANK1 | protein_coding | ENSG00000231863 | lncRNA | |
| ENSG00000297002 | | lncRNA | ENSG00000154655 | L3MBTL4 | protein_coding |
| ENSG00000150347 | ARID5B | protein_coding | ENSG00000144642 | RBMS3 | protein_coding |
| ENSG00000286339 | | lncRNA | ENSG00000136237 | RAPGEF5 | protein_coding |
| ENSG00000166510 | CCDC68 | protein_coding | ENSG00000164106 | SCRG1 | protein_coding |
| ENSG00000185386 | MAPK11 | protein_coding | ENSG00000306087 | lncRNA | |
| ENSG00000122641 | INHBA | protein_coding | ENSG00000308185 | lncRNA | |
| ENSG00000115738 | ID2 | protein_coding | ENSG00000235257 | ITGA9-AS1 | lncRNA |
| ENSG00000231925 | TAPBP | protein_coding | ENSG00000036565 | SLC18A1 | protein_coding |

| | | | | | |
|---|---|---|---|---|---|
| ENSG00000233871 | DLG5-AS1 | lncRNA | ENSG00000120160 | EQTN | protein_coding |
| ENSG00000136542 | GALNT5 | protein_coding | ENSG00000286615 | lncRNA | |
| ENSG00000144369 | FAM171B | protein_coding | ENSG00000138735 | PDE5A | protein_coding |
| ENSG00000245571 | FAM111A-DT | lncRNA | ENSG00000291034 | lncRNA | |
| ENSG00000169282 | KCNAB1 | protein_coding | ENSG00000018625 | ATP1A2 | protein_coding |
| ENSG00000232480 | TGFB2-AS1 | lncRNA | ENSG00000307088 | lncRNA | |
| ENSG00000188277 | C15orf62 | protein_coding | ENSG00000136943 | CTSV | protein_coding |
| ENSG00000135269 | TES | protein_coding | ENSG00000170417 | TMEM182 | protein_coding |
| ENSG00000137834 | SMAD6 | protein_coding | ENSG00000178297 | TMPRSS9 | protein_coding |
| ENSG00000203727 | SAMD5 | protein_coding | ENSG00000056998 | GYG2 | protein_coding |
| ENSG00000224843 | LINC00240 | lncRNA | ENSG00000251138 | LINC02882 | lncRNA |
| ENSG00000184497 | TMEM255B | protein_coding | ENSG00000169313 | P2RY12 | protein_coding |
| ENSG00000122786 | CALD1 | protein_coding | ENSG00000197299 | BLM | protein_coding |
| ENSG00000061455 | PRDM6 | protein_coding | ENSG00000175322 | ZNF519 | protein_coding |
| ENSG00000157110 | RBPMS | protein_coding | ENSG00000177910 | SPATA31C2 | protein_coding |
| ENSG00000263432 | RN7SL689P | misc_RNA | ENSG00000234511 | C5orf58 | protein_coding |
| ENSG00000135116 | HRK | protein_coding | ENSG00000175497 | DPP10 | protein_coding |
| ENSG00000278532 | | lncRNA | ENSG00000171408 | PDE7B | protein_coding |
| ENSG00000081320 | STK17B | protein_coding | ENSG00000225940 | C5orf67 | lncRNA |
| ENSG00000171246 | NPTX1 | protein_coding | ENSG00000145794 | MEGF10 | protein_coding |
| ENSG00000138821 | SLC39A8 | protein_coding | ENSG00000123901 | GPR83 | protein_coding |
| ENSG00000072858 | SIDT1 | protein_coding | ENSG00000232104 | RFX3-DT | lncRNA |
| ENSG00000288831 | | lncRNA | ENSG00000225889 | lncRNA | |
| ENSG00000240871 | KRTAP4-7 | protein_coding | ENSG00000008311 | AASS | protein_coding |
| ENSG00000232973 | CYP1B1-AS1 | lncRNA | ENSG00000230910 | ADGRB3-DT | lncRNA |
| ENSG00000172901 | LVRN | protein_coding | ENSG00000290931 | lncRNA | |
| ENSG00000166900 | STX3 | protein_coding | ENSG00000276672 | lncRNA | |

| | | | | | |
|---|---|---|---|---|---|
| ENSG00000225339 | | lncRNA | ENSG00000248441 | LETR1 | lncRNA |
| ENSG00000152804 | HHEX | protein_coding | ENSG00000171195 | MUC7 | protein_coding |
| ENSG00000125384 | PTGER2 | protein_coding | ENSG00000176714 | CCDC121 | protein_coding |
| ENSG00000069966 | GNB5 | protein_coding | ENSG00000205959 | lncRNA | |
| ENSG00000145808 | ADAMTS19 | protein_coding | ENSG00000223665 | lncRNA | |
| ENSG00000178538 | CA8 | protein_coding | ENSG00000228716 | DHFR | protein_coding |
| ENSG00000177822 | TENM3-AS1 | lncRNA | ENSG00000101695 | RNF125 | protein_coding |
| ENSG00000154127 | UBASH3B | protein_coding | ENSG00000163507 | CIP2A | protein_coding |
| ENSG00000286315 | | lncRNA | ENSG00000157851 | DPYSL5 | protein_coding |
| ENSG00000150637 | CD226 | protein_coding | ENSG00000148019 | CEP78 | protein_coding |
| ENSG00000072041 | SLC6A15 | protein_coding | ENSG00000239922 | lncRNA | |
| ENSG00000124788 | ATXN1 | protein_coding | ENSG00000198626 | RYR2 | protein_coding |
| ENSG00000178233 | TMEM151B | protein_coding | ENSG00000289212 | lncRNA | |
| ENSG00000245598 | DACT3-AS1 | lncRNA | ENSG00000129534 | MIS18BP1 | protein_coding |
| ENSG00000214106 | PAXIP1-AS2 | lncRNA | ENSG00000154920 | EME1 | protein_coding |
| ENSG00000263050 | DPH1-AS1 | lncRNA | ENSG00000129675 | ARHGEF6 | protein_coding |
| ENSG00000171502 | COL24A1 | protein_coding | ENSG00000235280 | MCF2L-AS1 | lncRNA |
| ENSG00000151067 | CACNA1C | protein_coding | ENSG00000150656 | CNDP1 | protein_coding |
| ENSG00000104450 | SPAG1 | protein_coding | ENSG00000156689 | GLYATL2 | protein_coding |
| ENSG00000187323 | DCC | protein_coding | ENSG00000140022 | STON2 | protein_coding |
| ENSG00000067955 | CBFB | protein_coding | ENSG00000257803 | PIGAP1 | processed_pseudogene |
| ENSG00000250508 | LINC02701 | lncRNA | ENSG00000272588 | lncRNA | |
| ENSG00000182511 | FES | protein_coding | ENSG00000118557 | PMFBP1 | protein_coding |
| ENSG00000228592 | | lncRNA | ENSG00000210195 | MT-TT | Mt_tRNA |
| ENSG00000307057 | | lncRNA | ENSG00000140525 | FANCI | protein_coding |
| ENSG00000120659 | TNFSF11 | protein_coding | ENSG00000287558 | lncRNA | |
| ENSG00000307789 | | lncRNA | ENSG00000305077 | lncRNA | |

| | | |
|---|---|---|
| ENSG00000249669 | CARMN | lncRNA |
| ENSG00000185650 | ZFP36L1 | protein_coding |
| ENSG00000177839 | PCDHB9 | protein_coding |
| ENSG00000286358 | | lncRNA |
| ENSG00000170959 | DCDC1 | protein_coding |
| ENSG00000229474 | PATL2 | protein_coding |
| ENSG00000180660 | MAB21L1 | protein_coding |
| ENSG00000172738 | TMEM217 | protein_coding |
| ENSG00000106034 | CPED1 | protein_coding |
| ENSG00000168386 | FILIP1L | protein_coding |
| ENSG00000188779 | SKOR1 | protein_coding |
| ENSG00000154783 | FGD5 | protein_coding |
| ENSG00000137573 | SULF1 | protein_coding |
| ENSG00000144724 | PTPRG | protein_coding |
| ENSG00000276740 | | lncRNA |
| ENSG00000164619 | BMPER | protein_coding |
| ENSG00000299656 | | lncRNA |
| ENSG00000184307 | ZDHHC23 | protein_coding |
| ENSG00000115267 | IFIH1 | protein_coding |
| ENSG00000287553 | | lncRNA |
| ENSG00000309889 | | lncRNA |
| ENSG00000137571 | SLCO5A1 | protein_coding |
| ENSG00000095585 | BLNK | protein_coding |
| ENSG00000125637 | PSD4 | protein_coding |
| ENSG00000105851 | PIK3CG | protein_coding |
| ENSG00000162804 | SNED1 | protein_coding |
| ENSG00000106785 | TRIM14 | protein_coding |
| ENSG00000172318 | B3GALT1 | protein_coding |
| ENSG00000169248 | CXCL11 | protein_coding |
| ENSG00000117724 | CENPF | protein_coding |
| ENSG00000113368 | LMNB1 | protein_coding |
| ENSG00000272841 | MAP3K4-AS1 | lncRNA |
| ENSG00000162374 | ELAVL4 | protein_coding |
| ENSG00000136457 | CHAD | protein_coding |

| | | |
|---|---|---|
| ENSG00000265972 | TXNIP | protein_coding |
| ENSG00000247809 | NR2F2-AS1 | lncRNA |
| ENSG00000126878 | AIF1L | protein_coding |
| ENSG00000069188 | SDK2 | protein_coding |
| ENSG00000294479 | | lncRNA |
| ENSG00000176595 | KBTBD11 | protein_coding |
| ENSG00000170961 | HAS2 | protein_coding |
| ENSG00000188610 | FAM72B | protein_coding |
| ENSG00000065923 | SLC9A7 | protein_coding |
| ENSG00000099957 | P2RX6 | protein_coding |
| ENSG00000204444 | APOM | protein_coding |
| ENSG00000100504 | PYGL | protein_coding |
| ENSG00000277135 | | lncRNA |
| ENSG00000183508 | TENT5C | protein_coding |
| ENSG00000255571 | MIR9-3HG | lncRNA |
| ENSG00000230294 | LINC02370 | lncRNA |
| ENSG00000110446 | SLC15A3 | protein_coding |
| ENSG00000137558 | PI15 | protein_coding |
| ENSG00000157734 | SNX22 | protein_coding |
| ENSG00000056998 | GYG2 | protein_coding |
| ENSG00000171843 | MLLT3 | protein_coding |
| ENSG00000259448 | LINC02352 | lncRNA |
| ENSG00000296522 | | lncRNA |
| ENSG00000226476 | LINC01748 | lncRNA |
| ENSG00000254689 | LINC02235 | lncRNA |
| ENSG00000198732 | SMOC1 | protein_coding |
| ENSG00000157510 | AFAP1L1 | protein_coding |

| | | |
|---|---|---|
| ENSG00000291034 | | lncRNA |
| ENSG00000132182 | NUP210 | protein_coding |
| ENSG00000238057 | ZEB2-AS1 | lncRNA |
| ENSG00000163449 | TMEM169 | protein_coding |
| ENSG00000171777 | RASGRP4 | protein_coding |
| ENSG00000144847 | IGSF11 | protein_coding |
| ENSG00000116962 | NID1 | protein_coding |
| ENSG00000174403 | CRMA | lncRNA |
| ENSG00000183798 | EMILIN3 | protein_coding |
| ENSG00000102445 | RUBCNL | protein_coding |
| ENSG00000253764 | KBTBD11-AS1 | lncRNA |
| ENSG00000279619 | | TEC |
| ENSG00000188783 | PRELP | protein_coding |
| ENSG00000251867 | | lncRNA |
| ENSG00000048740 | CELF2 | protein_coding |
| ENSG00000188176 | SMTNL2 | protein_coding |
| ENSG00000160862 | AZGP1 | protein_coding |
| ENSG00000132692 | BCAN | protein_coding |
| ENSG00000101198 | NKAIN4 | protein_coding |
| ENSG00000261329 | | lncRNA |
| ENSG00000248713 | C4orf54 | protein_coding |
| ENSG00000253931 | LINC02990 | lncRNA |
| ENSG00000188580 | NKAIN2 | protein_coding |
| ENSG00000064787 | BCAS1 | protein_coding |
| ENSG00000140450 | ARRDC4 | protein_coding |
| ENSG00000196091 | MYBPC1 | protein_coding |
| ENSG00000127084 | FGD3 | protein_coding |

| | | |
|---|---|---|
| ENSG00000122367 | LDB3 | protein_coding |
| ENSG00000145362 | ANK2 | protein_coding |
| ENSG00000122574 | WIPF3 | protein_coding |
| ENSG00000148200 | NR6A1 | protein_coding |
| ENSG00000100351 | GRAP2 | protein_coding |
| ENSG00000109062 | NHERF1 | protein_coding |
| ENSG00000144596 | GRIP2 | protein_coding |
| ENSG00000115525 | ST3GAL5 | protein_coding |
| ENSG00000101197 | BIRC7 | protein_coding |
| ENSG00000265142 | MIR133A1HG | lncRNA |
| ENSG00000280304 | | TEC |
| ENSG00000173210 | ABLIM3 | protein_coding |
| ENSG00000144868 | TMEM108 | protein_coding |
| ENSG00000141338 | ABCA8 | protein_coding |
| ENSG00000148082 | SHC3 | protein_coding |
| ENSG00000261121 | LINC02473 | lncRNA |
| ENSG00000225706 | PTPRD-AS1 | lncRNA |
| ENSG00000117069 | ST6GALNAC5 | protein_coding |
| ENSG00000162493 | PDPN | protein_coding |
| ENSG00000154914 | USP43 | protein_coding |
| ENSG00000172164 | SNTB1 | protein_coding |
| ENSG00000013619 | MAMLD1 | protein_coding |
| ENSG00000260947 | | lncRNA |
| ENSG00000260532 | | lncRNA |
| ENSG00000018625 | ATP1A2 | protein_coding |
| ENSG00000152932 | RAB3C | protein_coding |
| ENSG00000214548 | MEG3 | lncRNA |

| | | |
|---|---|---|
| ENSG00000104879 | CKM | protein_coding |
| ENSG00000120729 | MYOT | protein_coding |
| ENSG00000164649 | CDCA7L | protein_coding |
| ENSG00000278041 | | lncRNA |
| ENSG00000132688 | NES | protein_coding |
| ENSG00000108878 | CACNG1 | protein_coding |
| ENSG00000287792 | | lncRNA |
| ENSG00000175445 | LPL | protein_coding |
| ENSG00000158258 | CLSTN2 | protein_coding |
| ENSG00000235501 | CNN3-DT | lncRNA |
| ENSG00000145506 | NKD2 | protein_coding |
| ENSG00000047662 | FAM184B | protein_coding |
| ENSG00000187595 | ZNF385C | protein_coding |
| ENSG00000137393 | RNF144B | protein_coding |
| ENSG00000287385 | | lncRNA |
| ENSG00000248690 | HAS2-AS1 | lncRNA |
| ENSG00000125246 | CLYBL | protein_coding |
| ENSG00000248441 | LETR1 | lncRNA |
| ENSG00000221946 | FXYD7 | protein_coding |
| ENSG00000279622 | | lncRNA |
| ENSG00000243069 | ARHGEF26-AS1 | lncRNA |
| ENSG00000143195 | ILDR2 | protein_coding |
| ENSG00000170965 | PLAC1 | protein_coding |
| ENSG00000176402 | GJC3 | protein_coding |
| ENSG00000227082 | LINC02798 | lncRNA |
| ENSG00000197106 | SLC6A17 | protein_coding |
| ENSG00000005981 | ASB4 | protein_coding |

| | | |
|---|---|---|
| ENSG00000164946 | FREM1 | protein_coding |
| ENSG00000183230 | CTNNA3 | protein_coding |
| ENSG00000142661 | MYOM3 | protein_coding |
| ENSG00000287161 | | lncRNA |
| ENSG00000129682 | FGF13 | protein_coding |
| ENSG00000239922 | | lncRNA |
| ENSG00000164879 | CA3 | protein_coding |
| ENSG00000170417 | TMEM182 | protein_coding |
| ENSG00000168843 | FSTL5 | protein_coding |
| ENSG00000147869 | CER1 | protein_coding |
| ENSG00000163406 | SLC15A2 | protein_coding |
| ENSG00000131730 | CKMT2 | protein_coding |
| ENSG00000254656 | RTL1 | protein_coding |
| ENSG00000254338 | MAFA-AS1 | lncRNA |
| ENSG00000305242 | | lncRNA |
| ENSG00000159399 | HK2 | protein_coding |
| ENSG00000224945 | | lncRNA |
| ENSG00000275620 | | lncRNA |
| ENSG00000114529 | C3orf52 | protein_coding |
| ENSG00000174611 | KY | protein_coding |
| ENSG00000124507 | PACSIN1 | protein_coding |
| ENSG00000133256 | PDE6B | protein_coding |
| ENSG00000138193 | PLCE1 | protein_coding |
| ENSG00000115592 | PRKAG3 | protein_coding |
| ENSG00000260442 | ATP2A1-AS1 | lncRNA |
| ENSG00000109063 | MYH3 | protein_coding |
| ENSG00000187699 | C2orf88 | protein_coding |

| | | |
|---|---|---|
| ENSG00000234511 | C5orf58 | protein_coding |
| ENSG00000155886 | SLC24A2 | protein_coding |
| ENSG00000117115 | PADI2 | protein_coding |
| ENSG00000101680 | LAMA1 | protein_coding |
| ENSG00000287781 | | lncRNA |
| ENSG00000104490 | NCALD | protein_coding |
| ENSG00000136842 | TMOD1 | protein_coding |
| ENSG00000183873 | SCN5A | protein_coding |
| ENSG00000229891 | LINC01315 | lncRNA |
| ENSG00000291233 | | lncRNA |
| ENSG00000231473 | RB1-DT | lncRNA |
| ENSG00000205929 | EPCIP | protein_coding |
| ENSG00000170379 | TCAF2 | protein_coding |
| ENSG00000175567 | UCP2 | protein_coding |
| ENSG00000101203 | COL20A1 | protein_coding |
| ENSG00000230498 | | lncRNA |
| ENSG00000130595 | TNNT3 | protein_coding |
| ENSG00000238906 | | snoRNA |
| ENSG00000180354 | MTURN | protein_coding |
| ENSG00000239828 | CCDC54-AS1 | lncRNA |
| ENSG00000260398 | | lncRNA |
| ENSG00000177807 | KCNJ10 | protein_coding |
| ENSG00000185668 | POU3F1 | protein_coding |
| ENSG00000171033 | PKIA | protein_coding |
| ENSG00000231698 | | lncRNA |
| ENSG00000138944 | SHISAL1 | protein_coding |
| ENSG00000106772 | PRUNE2 | protein_coding |

| | | |
|---|---|---|
| ENSG00000127252 | PLAAT1 | protein_coding |
| ENSG00000285671 | | lncRNA |
| ENSG00000270071 | | lncRNA |
| ENSG00000135636 | DYSF | protein_coding |
| ENSG00000070182 | SPTB | protein_coding |
| ENSG00000103994 | ZNF106 | protein_coding |
| ENSG00000108405 | P2RX1 | protein_coding |
| ENSG00000287063 | | lncRNA |
| ENSG00000124839 | RAB17 | protein_coding |
| ENSG00000086967 | MYBPC2 | protein_coding |
| ENSG00000137124 | ALDH1B1 | protein_coding |
| ENSG00000101204 | CHRNA4 | protein_coding |
| ENSG00000116396 | KCNC4 | protein_coding |
| ENSG00000036448 | MYOM2 | protein_coding |
| ENSG00000184343 | SRPK3 | protein_coding |
| ENSG00000154556 | SORBS2 | protein_coding |
| ENSG00000133020 | MYH8 | protein_coding |
| ENSG00000144485 | HES6 | protein_coding |
| ENSG00000250421 | | lncRNA |
| ENSG00000214114 | MYCBP | protein_coding |
| ENSG00000134207 | SYT6 | protein_coding |
| ENSG00000131725 | WDR44 | protein_coding |
| ENSG00000166257 | SCN3B | protein_coding |
| ENSG00000281128 | PTENP1-AS | lncRNA |
| ENSG00000170807 | LMOD2 | protein_coding |
| ENSG00000158296 | SLC13A3 | protein_coding |
| ENSG00000290923 | PGM5P2 | lncRNA |

| | | |
|---|---|---|
| ENSG00000164867 | NOS3 | protein_coding |
| ENSG00000172403 | SYNPO2 | protein_coding |
| ENSG00000185739 | SRL | protein_coding |
| ENSG00000136546 | SCN7A | protein_coding |
| ENSG00000286924 | | lncRNA |
| ENSG00000165996 | HACD1 | protein_coding |
| ENSG00000263105 | | lncRNA |
| ENSG00000143632 | ACTA1 | protein_coding |
| ENSG00000185567 | AHNAK2 | protein_coding |
| ENSG00000115593 | SMYD1 | protein_coding |
| ENSG00000057704 | TMCC3 | protein_coding |
| ENSG00000181085 | MAPK15 | protein_coding |
| ENSG00000053918 | KCNQ1 | protein_coding |
| ENSG00000125414 | MYH2 | protein_coding |
| ENSG00000301470 | | lncRNA |
| ENSG00000304025 | | lncRNA |
| ENSG00000171766 | GATM | protein_coding |
| ENSG00000163297 | ANTXR2 | protein_coding |
| ENSG00000083454 | P2RX5 | protein_coding |
| ENSG00000140057 | AK7 | protein_coding |
| ENSG00000172508 | CARNS1 | protein_coding |
| ENSG00000197361 | FBXL22 | protein_coding |
| ENSG00000171234 | UGT2B7 | protein_coding |
| ENSG00000272927 | | lncRNA |
| ENSG00000102678 | FGF9 | protein_coding |
| ENSG00000172000 | ZNF556 | protein_coding |
| ENSG00000310298 | | lncRNA |

| | | |
|---|---|---|
| ENSG00000153531 | ADPRHL1 | protein_coding |
| ENSG00000151952 | TMEM132D | protein_coding |
| ENSG00000203867 | RBM20 | protein_coding |
| ENSG00000130433 | CACNG6 | protein_coding |
| ENSG00000250899 | | lncRNA |
| ENSG00000172995 | ARPP21 | protein_coding |
| ENSG00000197769 | MAP1LC3C | protein_coding |
| ENSG00000261553 | | lncRNA |
| ENSG00000070808 | CAMK2A | protein_coding |
| ENSG00000081248 | CACNA1S | protein_coding |
| ENSG00000196169 | KIF19 | protein_coding |

Mt_tRNA stands for mitochondrial transfer RNA, lncRNA encodes for long non-coding RNA, misc_RNA is short for miscellaneous RNA, TEC stands for To be Experimentally Confirmed.

**Supplementary Table S5.** List of antibodies

| Antibody | Supplier and reference | Dilution |
|---|---|---|
| Goat IgG anti-NANOG | AF1997, R&D Systems | 1:25 |
| Mouse IgM anti-TRA-1-81 | MAB4381, Millipore | 1:400 |
| Mouse IgG anti-OCT4 | 60059, Stem Cell Technologies | 1:100 |
| Rat IgM anti-SSEA3 | MC-631, Hybridoma Bank | 1:3 |
| Rabbit IgG anti-SOX2 | PA1-16968, Pierce Antibodies | 1:100 |
| Mouse IgG anti-SSEA4 | MC-813-70, Hybridoma Bank | 1:3 |
| Mouse IgM anti-Tra-1-60 | MAB4360, Sigma, Aldrich | 1:400 |
| Mouse IgG anti-ASMA | A5228, Sigma | 1:400 |
| Mouse IgM anti-ASA | A2172, Sigma | 1:400 |
| Rabbit IgG anti-GATA4 | Sc-9053, Santa Cruz Biotechnology | 1:50 |
| Mouse IgG anti-AFP | MAB1368, R&D Systems | 1:50 |
| Goat IgG anti-FOXA2 | AF2400, R&D Systems | 1:50 |
| Mouse IgG anti-TUJ1 | MMS-435P, BioLegend | 1:500 |
| Rabbit IgG anti GFAP | Z0334, Dako | 1:500 |
| Rabbit IgG anti-Vinculin | ab109244, Abcam | 1:500 |
| Rabbit IgG anti-merlin | Ab109244, Abcam | 1:5000 |
| Mouse IgG anti [NGFR5] to p75 NGF Receptor | ab3125, Abcam | 1:100 (IF)<br>1:1000 (FACS) |
| Rabbit IgG anti-S100B | Z0311, Dako | 1:1000 |
| Rabbit IgG anti-Sox10 | ab155279, Abcam | 1:50 |
| Mouse IgG anti-AP2 | MA1-872, Thermo Scientific | 1:50 |
| Mouse IgG anti-HNK1 | C6680, Sigma | 1:1000 |
| Rabbit IgG anti-pS6 | 2211L, Cell Signaling Technology | 1:1000 |
| Rabbit IgG anti-S6 | 2217S, Cell Signaling Technology | 1:1000 |
| Mouse IgG-pAkt | 4051S, Cell Signaling Technology | 1:1000 |
| Rabbit IgG anti-Akt | 9272S, Cell Signaling Technology | 1:1000 |
| Rabbit IgG anti-Cyclin-D1 | 55506T, Cell Signaling Technology | 1:1000 |
| Mouse IgG anti-YAP1(H-9) | sc-271134, Santa Crus Biotechnology | 1:100 |

**Supplementary Table S6.** *In vivo* Phosphorodiamidate Morpholino Oligomers sequences targeting exon 11 of *NF2* gene

| ***In vivo* Phosphorodiamidate Morpholino Oligomers ES11** | |
|---|---|
| **PMO 5’ sequence** | CGCTCCATCTGCGAGGGGTGAAGAA |
| **PMO 3’ sequence** | CCTCAGAAATCACCAGTGCTTCGTT |

### Supplementary Figures

**Fig S1: *NF2* cDNA gene characterization after CRISPR/Cas9 editing.**
**A:** Sanger sequencing of the generated and selected clones, LOF variants were sequenced directly from the iPSC cell culture. Reverse sequence of exon 11 of *NF2* gene is shown. **B:** SNP-array analysis showed no loss of heterozygosity (LOH) and no presence of complex rearrangements. *NF2*(+/-) and *NF2*(-/-) lines showed no differences with respect to the line of origin, *NF2*(+/+). BAF: B allele frequency; LRR: Log R Ratio.

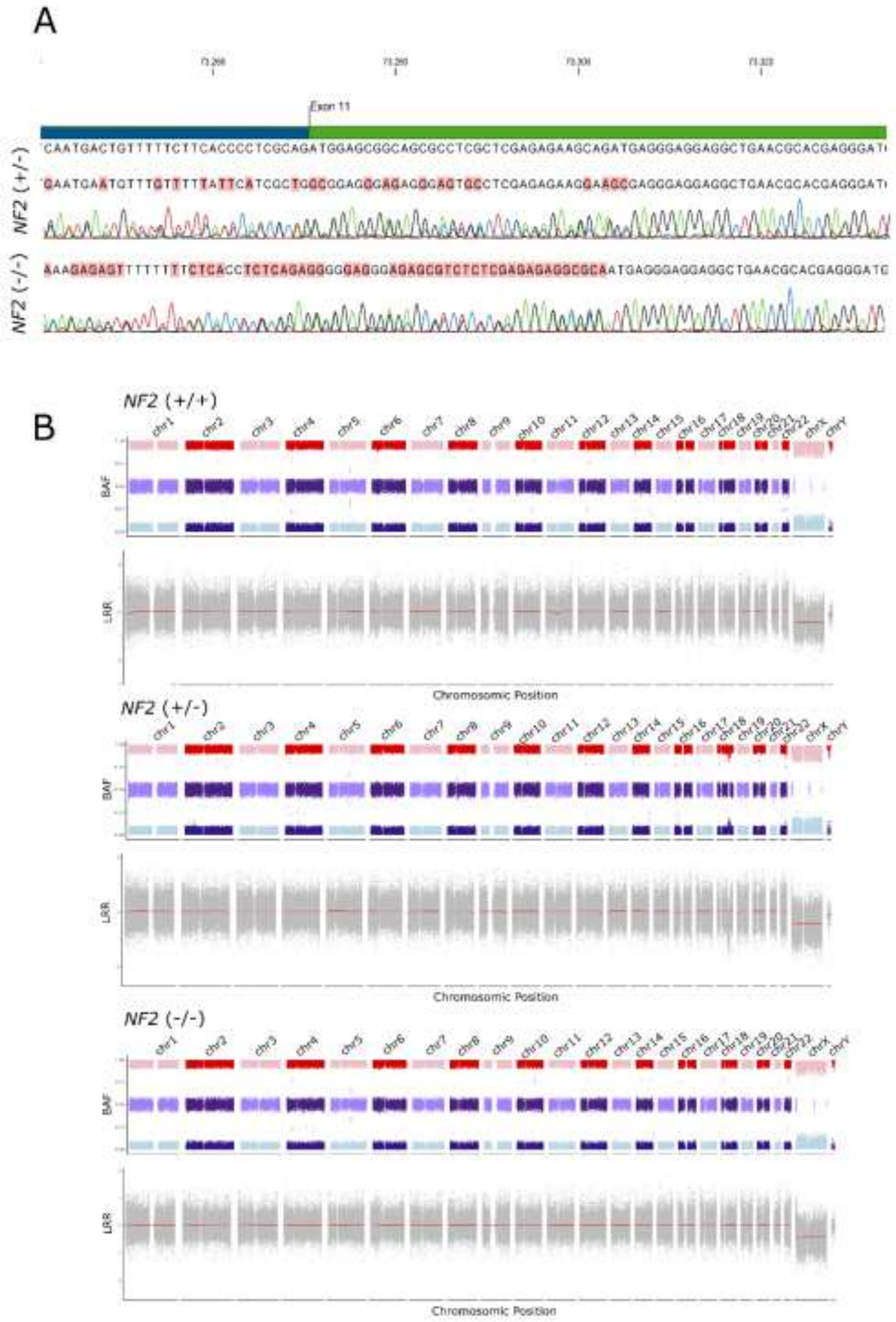

**Fig S2. Expression profile of *NF2*(+/-) and *NF2*(-/-) generated lines**

**A:** Principal Component Assay (PCA) shows obtained cell lines speared by genotype (*NF2*(+/+), *NF2*(+/-) and *NF2*(-/-)) and cell type (iPSCs, NC, SC differentiated for 7 days [SC7] and SC differentiated for 14 days [SC14]). **B:** GSEA of *NF2*(+/-) or *NF2*(-/-) and *NF2*(+/+) Schwann Cells differentiated for 14 days respectively. NES stands for Normalized Enrichment Score. Upregulated pathways are represented in blue, downregulated pathways represented in red. Results are filtered by p value (<0.05). **C:** Differential expression analysis of *NF2*-related pathways by Single Sample GSEA comparison. Genotype stands for *NF2*(+/+), *NF2*(+/-) and *NF2*(-/-) respectively. TF means Transcription Factor. Lines represent the mean enrichment score of every genotype.

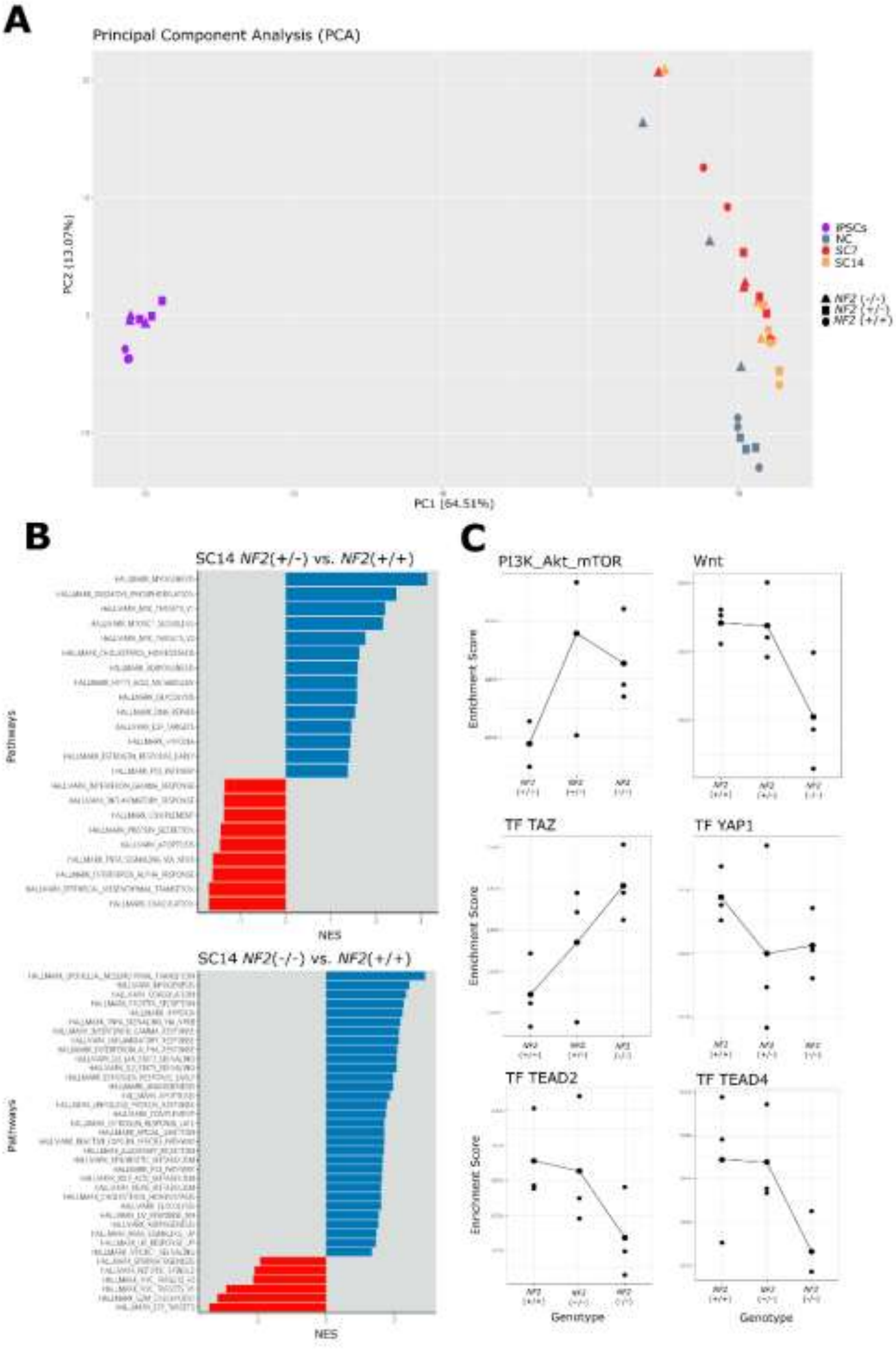

**Fig S3 Engraftment of Schwann Cell spheroids on nude mice**

**A:** Schematic representation of the experimental design applied for the engraftment the SC spheroids in nude mouse. **B:** Hematoxylin and eosin morphological characterization of the gastrocnemius muscle of the engrafted mice. Scale bar: 100 µm.

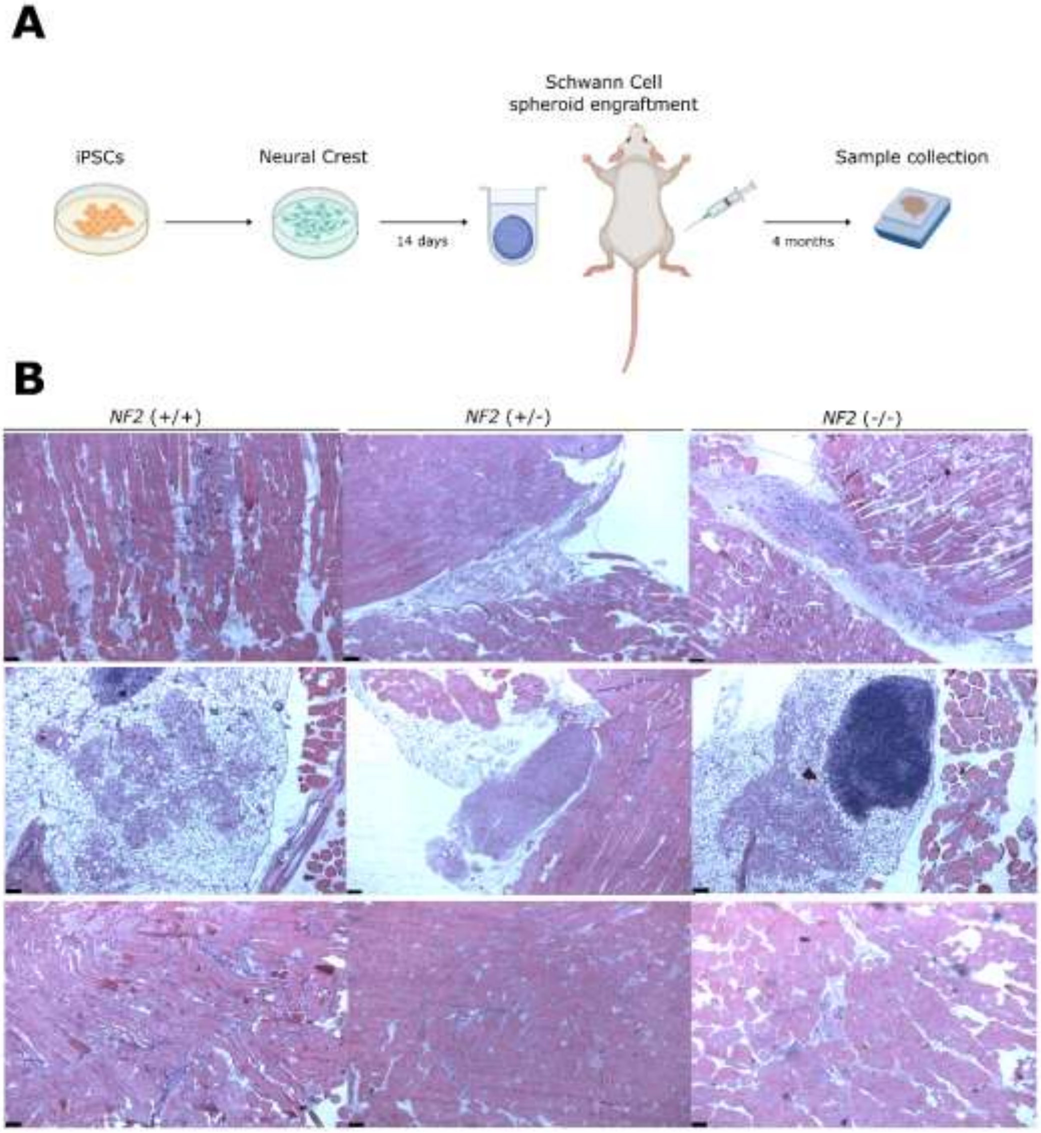

**Fig S4 Unspecific effect of PMO Treatment**

**A:** Volcano plot shows comparison of *NF2*(+/-) SC spheroids after 7 days of treatment (T) and without treatment (UT) [top]. Same comparison was assessed for *NF2*(-/-) SC spheroids [bottom]. Significantly upregulated and downregulated genes are displayed in green and red, respectively. Blue dots stand for DEGs related to unspecific effect of PMO treatment. Significantly expressed genes are those with an absolute change of expression > 1 and adjusted p value < 0.01.

**A**

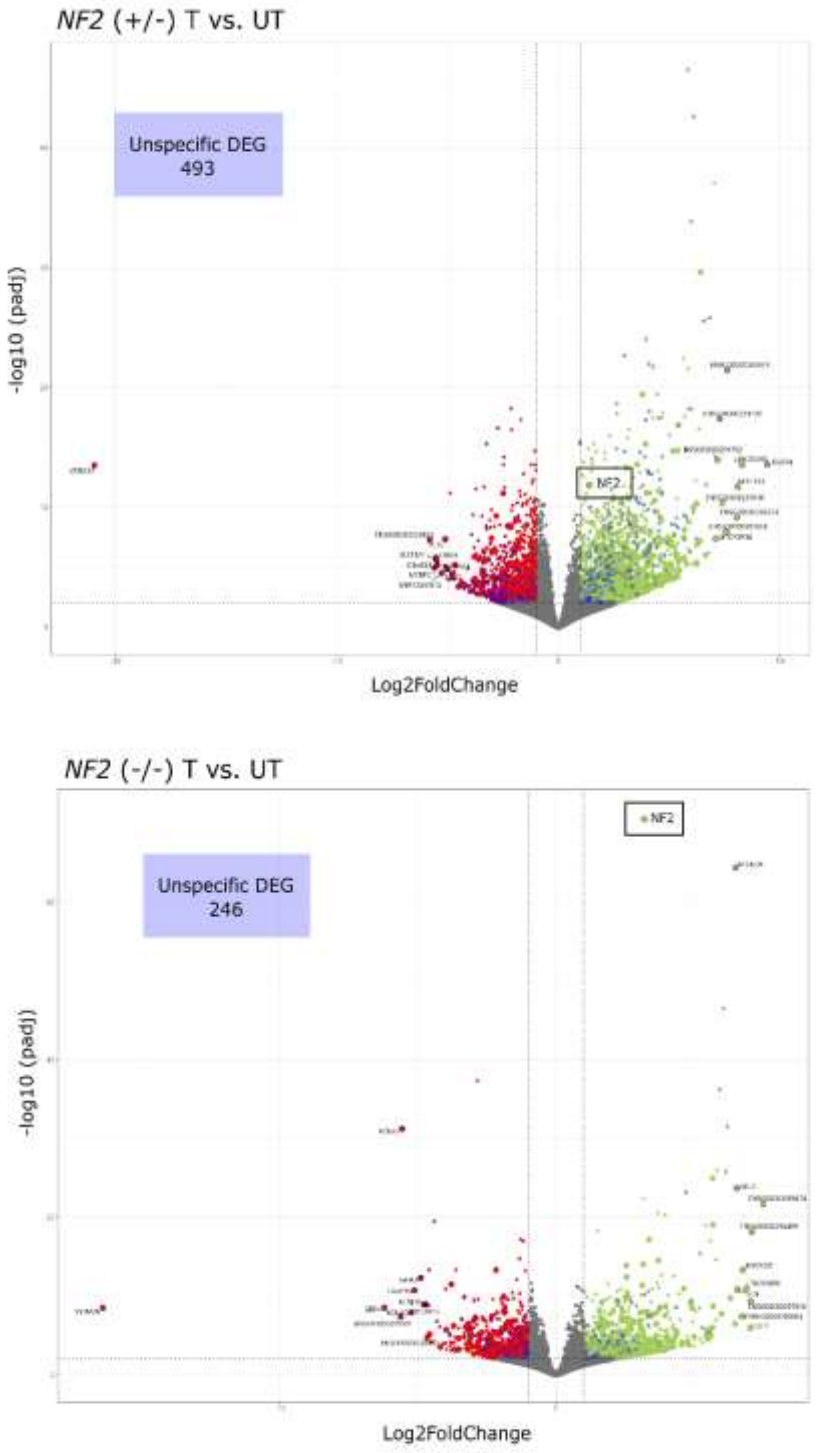

**Fig S5 String of the unspecific effect of PMO Treatment**

**A:** STRING analysis was assessed for all DEGs related to the unspecific effect of PMO treatment. DEGs with an interaction score > 0.7000 are shown. *NF2* is highlighted in red.

**A1** *NF2* (+/-)

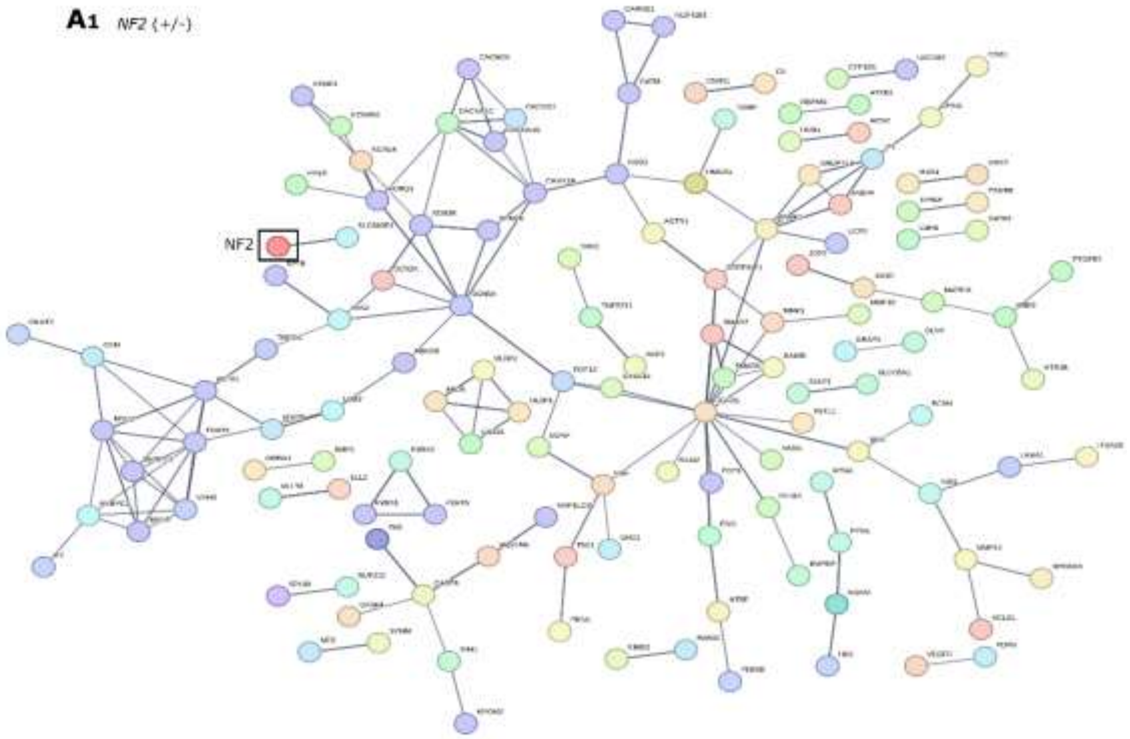


**A2** *NF2* (-/-)

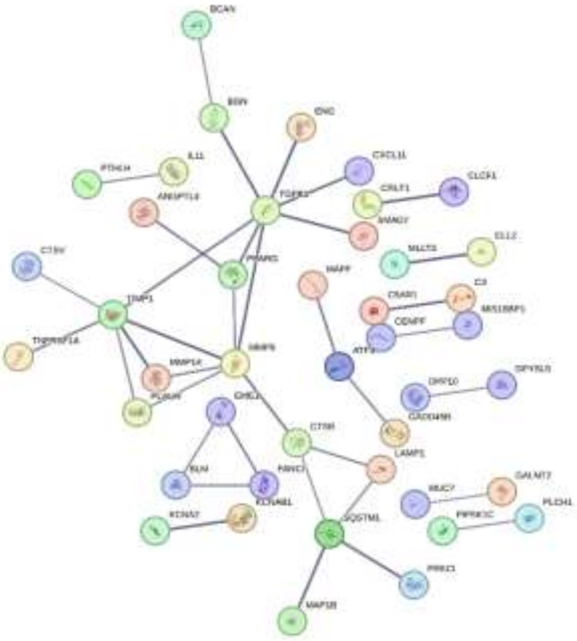

**Fig S6 Enrichment analysis of Schwann Cell spheroids after treatment**

**A:** Enrichment analysis was assessed for SC spheroids after 7 days of PMO treatment. GO Biological Process is shown. A1: *NF2* (+/-) upregulated pathways. A2: *NF2*(+/-) downregulated pathways. A3: *NF2*(-/-) upregulated pathways. A4: *NF2*(-/-) downregulated pathways. The shown pathways are the top 15 enriched at the FDR level and are ordered by count.

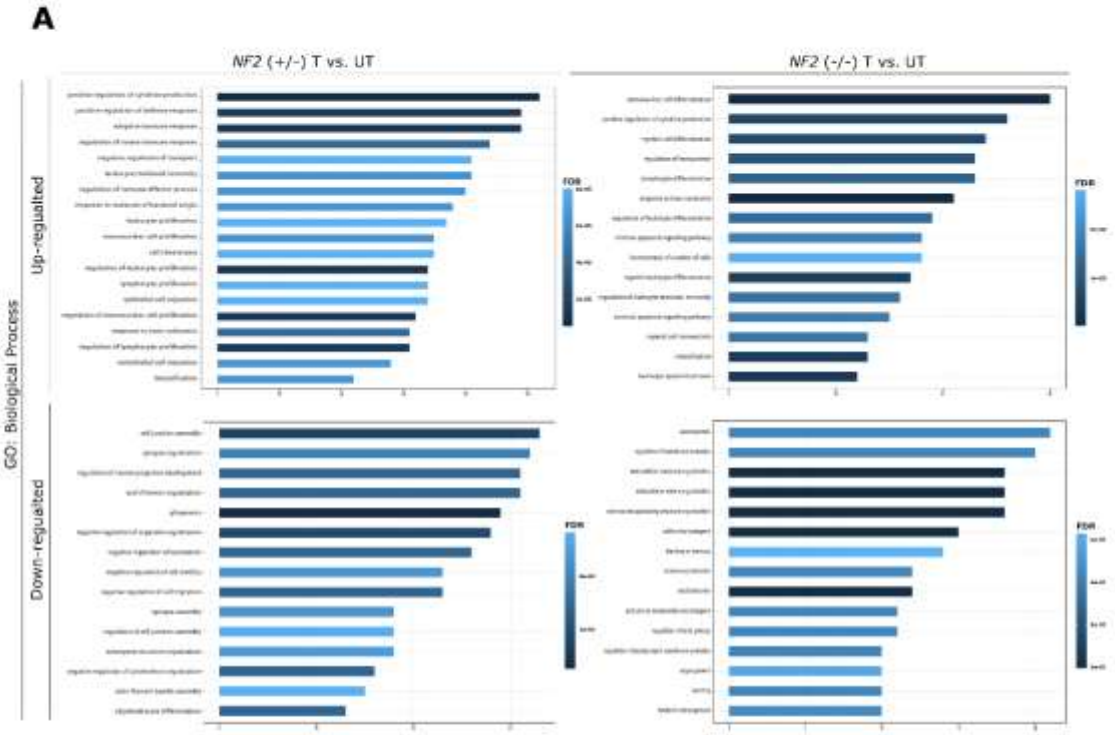
